# Simultaneous discovery of candidate imprinted genes and Imprinting Control Regions in the mouse genome

**DOI:** 10.1101/780551

**Authors:** Minou Bina, Phillip Wyss

## Abstract

In mammals, parent-of-origin-specific gene expression is regulated by specific genomic DNA segments known as Imprinting Control Regions (ICRs) and germline Differentially Methylated Regions (gDMRs). In the mouse genome, the known ICRs/gDMRs often include clusters of a set of composite-DNA-elements known as ZFBS-morph overlaps. These elements consist of the ZFP57 binding site (ZFBS) overlapping a subset of the MLL1 morphemes. To improve detection of such clusters, we created density-plots. In genome-wide analyses, peaks in these plots pinpointed ∼90% of the known ICRs/gDMRs and located candidate ICRs within relatively long genomic DNA sections. In several cases, the candidate ICRs mapped to chromatin boundaries, to a subset of gene-transcripts, or to both. By viewing the plots at the UCSC genome browser, we could examine the candidate ICRs in the context of the genes in their vicinity. This strategy uncovered several potential imprinted genes with a broad range of physiologically important functions. Examples include: folliculogenesis; lineage commitment of murine embryonic stem cells; the development of the junctional zone of the placenta; left-right patterning of the body axis; the development of the neocortex, hippocampus, and cerebellum; postnatal vision; self-renewal of mouse spermatogonial stem cells; and histone-to-protamine replacement during spermatogenesis.

## INTRODUCTION

In mammals, the inherited pairs of chromosomes are not functionally identical [1-3]. In the course of gametogenesis, genomic imprinting affects a subset of genes and results in monoallelic, parental-of-origin specific expression [4, 5]. The imprinting process is relatively complex. It involves chromatin remodeling, methylation of the ICRs/DMRs in one the parental alleles, and binding of ZFP57 to a subset of the modified CpGs to maintain the DNA methylation memory [6]. Among the DNA methylation systems, a protein-complex consisting of DNMT3A and DNMT3L plays a central role in genomic imprinting [6, 7]. This complex methylates DNA processivity [8]. The rate of product formation is influenced by the number of CpGs, particularly by those dispersed in the CpG islands (CGIs) associated with gene promoters [8]. Within these islands are clusters of CpG-rich motifs known as the MLL1 morphemes [9]. The MT domain in MLL1 and a related family member (MLL2) binds CpG-rich motifs [10, 11]. Both MLL1 and MLL2 function in promoter-specific trimethylation of histone H3 lysine 4 producing H3K4me3 marks in chromatin [12]. Besides gene promoters, the CpG islands also occur within intergenic and intragenic sequences [9, 13]. Promoter-associated islands often extend into the transcribed sequences, even into protein-coding exons [14]. The occurrences of the MLL1 morphemes in these exons impact codon utilization and preservation [15], perhaps reflecting the importance of these CpG-rich motifs in genome-associated functions.

The fully characterized ICRs/gDMRs often include sequences known as ZFBS-morph overlaps [13, 16]. These composite-DNA-elements are CpG-rich and thus could facilitate the processive methylation of several CpGs by the DNMT3A-DNMT3L complex [16]. While *in vitro* ZFP57 binds methylated DNA [17], the MT domain in MLL1 and MLL2 selectively interacts with unmodified sequences [10, 11]. Overall, CpG dinucleotides occur infrequently in animal DNA [18]. Thus, in mammalian DNA the hexameric ZFP57 binding site occurs less often than sequences that do not contain any CpG [17]. Moreover, since the composite DNA elements contain 2 or more CpGs, their occurrences in the DNA are far less than those observed for the canonical ZFP57 binding site [16]. Since clusters of 2 or more ZFBS-morph overlaps located ∼ 90% of the characterized ICRs/gDMRs in the mouse genome [17], we wished to explore whether locating such clusters could pinpoint candidate ICRs and reveal the genomic positions of potential imprinted genes. However, even though the ZFBS-morph overlaps represent only ∼ 12.7% of ZFP57 binding sites along Chr7 [16], their isolated occurrences create background noise. A noisy background would interfere with discerning the genomic positions of the ICRs/gDMRs within relatively long DNA sections. To resolve this problem, we devised a bioinformatics strategy. The scheme consisted of creating plots to display the density of ZFBS-morph overlaps in mouse genomic DNA. Previously, this approach facilitated locating clusters of ZFBS-morph overlaps in the human genome [19]. To explore the predictive capacity of our strategy for the mouse genome, we sampled the ‘robust’ peaks in the density-plots, to identify the genes or transcripts in their vicinity. The examined samples mapped to genes important to fetal growth, vision, organogenesis, neurogenesis, and sex-related processes.

## 2. RESULTS

We wrote a Perl script that employed a specified sliding window to determine and to report the density of ZFBS-Morph overlaps along chromosomal DNA sequences. We chose the window-size (the number of covered DNA bases) by trial and error. To do so, we selected the extensively characterized ICR of the *H19* – *Igf2* imprinted domain to gauge the appearance of peaks in the density-plots. We noticed that long windows gave spurious peaks; small windows had a spikey appearance. Based on this and related exploratory studies, we chose a window consisting of 850 bases. For this window-size, the ICR of the *H19* - *Igf2* imprinted domain appears under two density peaks (Fig. 1). This ICR includes several chromatin boundaries [20, 21], and two ZFP57 associated regions [22]. The locations of the two density peaks agree with positions of ZFP57associated regions and the chromatin boundaries within the ICR of the *H19* - *Igf2* imprinted domain.

**Figure 1.**
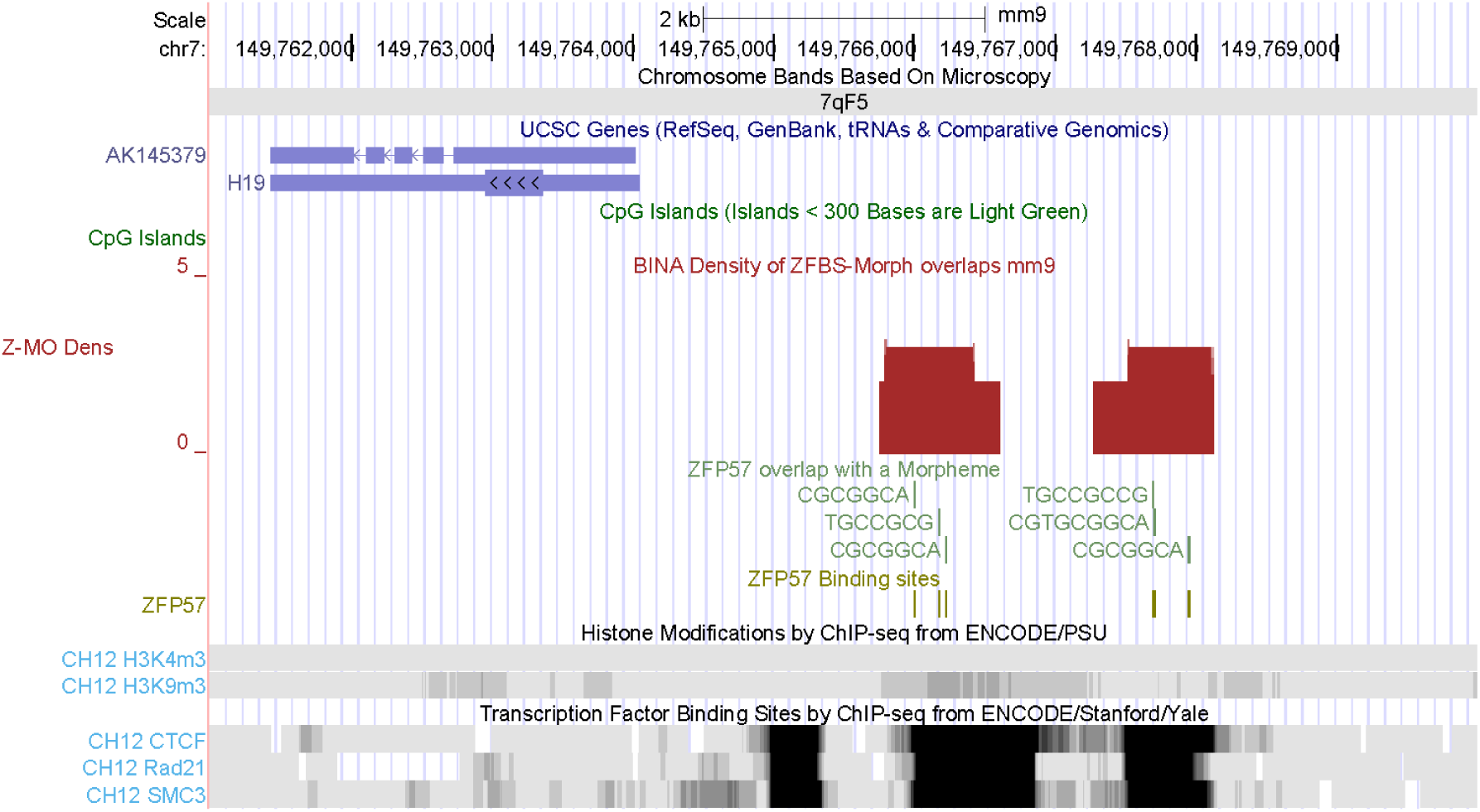
Genomic position of the ICR in the *H19* – *Igf2* imprinted domain in the 850-base window selected for creating the density-plots. Note that the ICR does not include H3K9me3 marks. This is because with ChIPs, occasionally the antibody cannot detect highly condensed regions in chromatin.

In preliminary assessments, we found that density peaks covering 2 ZFBS-Morph overlaps could be false or true-positive. Therefore, to locate candidate ICRs, we focused on the ‘robust’ peaks encompassing 3 or more ZFBS-Morph overlaps. In initial evaluations, we asked whether we could spot the known ICRs/gDMRs within relatively long genomic DNA sections. Concurrently, we inspected these DNA sections for peaks that may reflect the genomic positions of candidate ICRs. By displaying the density-plots at the UCSC browser, we could obtain close-up views to investigate the positions of the candidate ICRs with respect to genomic landmarks; including genes, transcripts, and the CpG islands. We also sampled randomly selected DNA segments to determine whether they included chromatin boundaries, genes important to embryonic development, or both.

### Locating the position of ICRs/gDMRs in relatively long genomic DNA sections

#### 1. Density peaks revealed the essential ICR/gDMR of the complex Gnas locus and candidate ICRs within the Lsm14b and Taf4a loci

The *Gnas* complex locus is in Chr2qH4. This chromosomal band contains ∼ 8.25 Mb DNA (Fig. 2). Along the displayed segment, discernable are 3 robust density peaks and several peaks covering 2 ZFBS-Morph overlaps. Thus, for that DNA section we observed 1 robust peak per ∼ 2.57 Mb DNA. One of the 3 robust peaks located the essential ICR of the complex *Gnas* locus. The other 2 correspond to candidate ICRs (Fig. 2). In enlarged views, the peak in the *Gnas* locus appeared as a doublet defining the two distinguishable clusters of ZFBS-Morph overlaps in the locus [13, 16]. In the DNA, a candidate ICRs maps to *Taf4a* (a subunit of the general transcription factor TFIID). Another one maps to *Lsm14b* (LSM family member 14B). From *Lsm14b* are produced three transcripts with differing exon utilization patterns (Fig. S1). The density peak is in the CGI that encompasses the promoter and the TSS of the longest *Lsm14b* transcript (Fig. S1). Functionally, LSM14B is an RNA binding protein that plays an essential role in oocyte meiotic maturation through regulating mRNA pools [23]. For *Taf4a*, genomic maps also revealed several transcripts. The candidate ICR maps to the 1^st^ exon of the longest *Taf4a* transcript (Fig. S1). The corresponding density peak is between two CpG islands. One of the islands encompasses a bidirectional-promoter regulating expression from *Taf4a* and *4921531C22Rik*. According to GENE at NCBI, *4921531C22Rik* is broadly expressed in adult testes (https://www.ncbi.nlm.nih.gov/gene/66730#gene-expression). Since the TSSs of *Taf4a* and *4921531C22Rik* are not far-apart (Fig. S1), the candidate ICR may regulate allele-specific expression of both *4921531C22Rik* and the longest *Taf4a* transcript.

**Figure 2.**
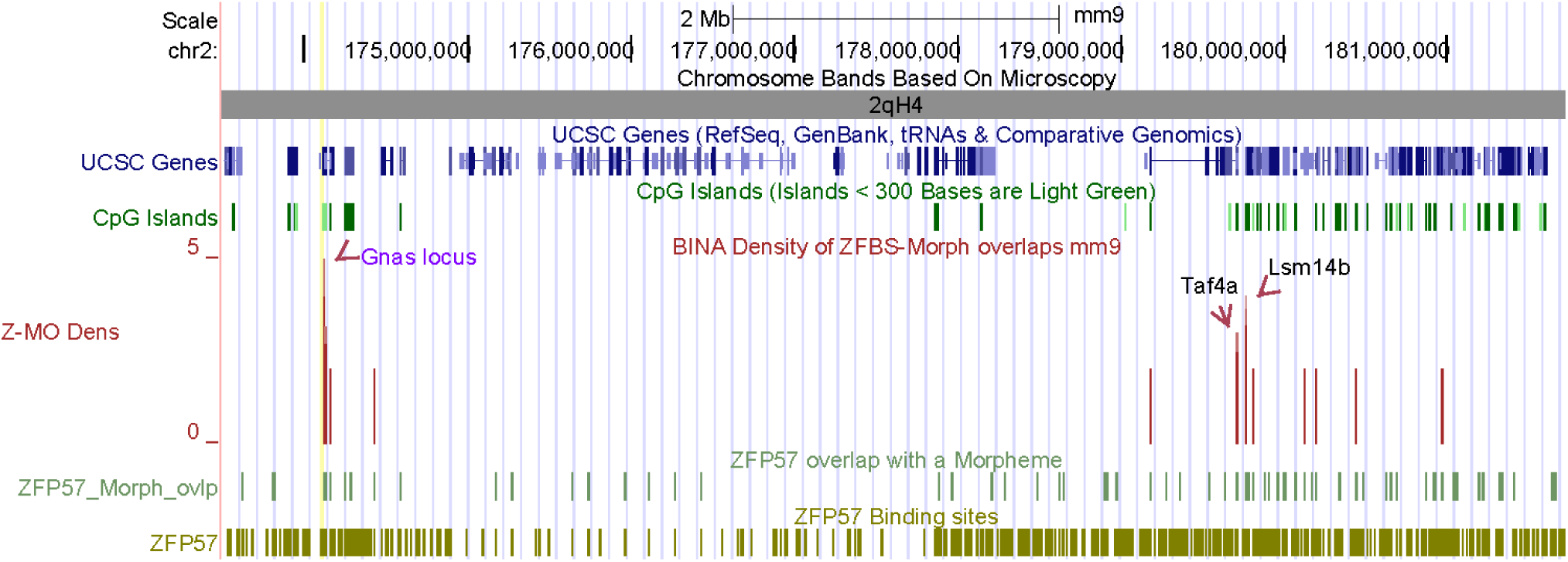
Genomic positions of density peaks along Chr2qH4. The displayed tracks give the locations of chromosomal bands (gray), genes (blue), the CpG islands (green), density peaks (maroon), the ZFBS-morph overlaps (hunter green), and the ZFP57 binding sites (olive). The density-plot is shown in full format. The remaining tracks are shown in dense format. An arrow points to the peak corresponding to the essential ICR of the *Gnas* complex locus (shown in purple). Two arrows point to candidate ICRs near the *Taf4a* and *Lsm14b* loci. Note that at such a low-resolution physical map (covering ∼ 8.25 Mb DNA), clearly resolved are density peaks covering 2 or more ZFBS-morph overlaps. In contrast, the positions of the CpG islands are not resolved. The occurrences of the hexameric ZFP57 site are not clearly apparent. Even though less frequent, there are also many isolated ZFBS-morph overlaps.

#### 2. Density peaks pinpointed the ICR/gDMR of Mest and revealed candidate ICRs for potential imprinted genes

The ICR of *Mest* is in Chr6qA. This chromosomal band consists of several subbands. The entire band encompasses nearly 35 Mb long DNA and 4 robust density peaks (Fig. 3). Thus, for that DNA section we observed 1 robust peak per ∼ 8.7 Mb DNA. We noticed that overall, robust peaks were not uniformly distributed along the chromosomal DNA sequences. For example, 4 consecutive subbands (6qA1, 6qA2, 6qA3.1, 6qA3.2) contain only peaks covering 2 ZFBS-Morph-overlaps. In contrast, 6qA3.3 encompasses the 4 robust density peaks (Fig. 3). One of the peaks spotted the known ICR/gDMR in the *Mest* locus. The other 3 correspond to candidate ICRs. They map to: (i) a CGI that encompasses the TSS of *Impdh1* (Fig. S2); (ii) a CGI that that encompasses the TSS of *Fam40b* (Fig. S3); and (iii) a region far upstream of the TSS of *Slc35b4* (Fig. S4).

**Figure 3.**
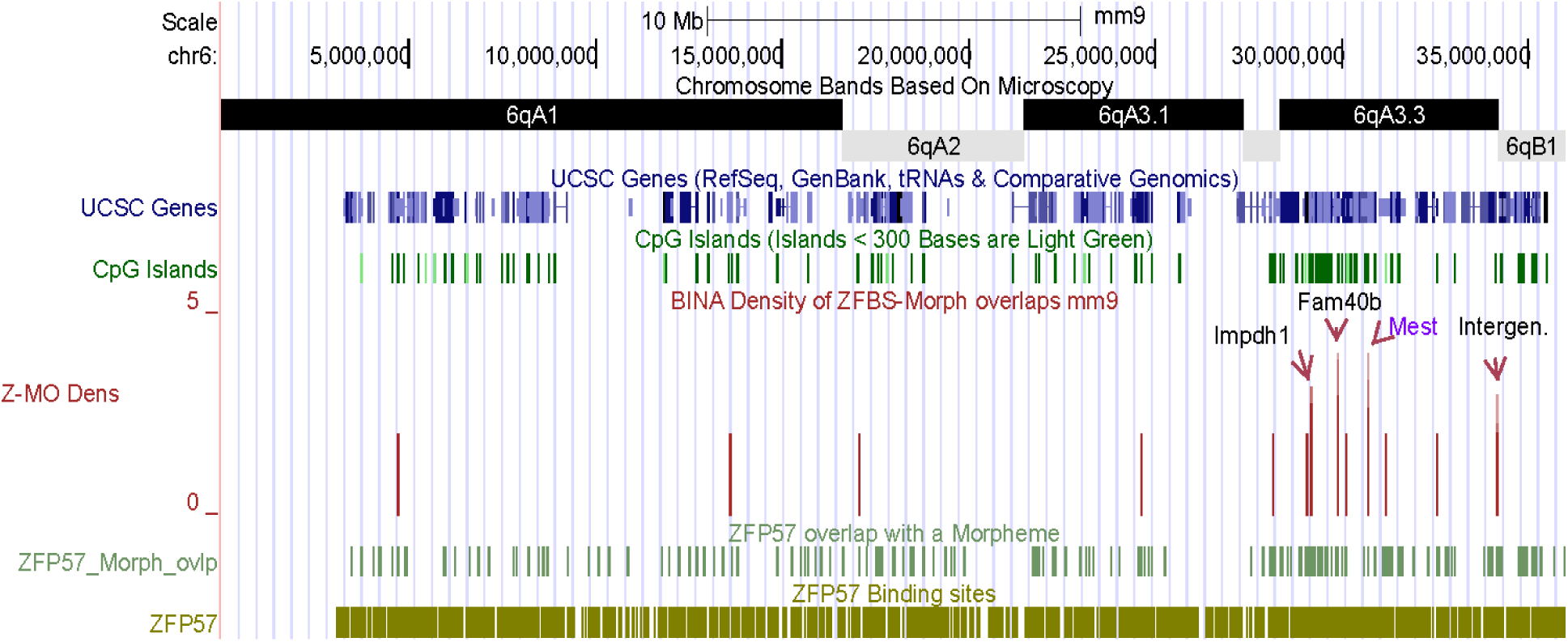
Genomic positions of robust density peaks along Chr6qA. One peak spotted the known ICR/gDMR of the *Mes*t locus. Several peaks mark the position of candidate ICRs regulating expression of potential imprinted genes.

Functionally, IMPDH1 (inosine-5’-monophosphate dehydrogenase 1) catalyzes the rate-limiting step in biosynthesis of guanine nucleotides. Mutations in the corresponding human gene caused retinitis pigmentosa, an inherited retinal degeneration characterized by the early onset of night blindness followed by a progressive loss of the visual field [24, 25]. Also known as *Strip2* (striatin interacting protein 2), *Fam40b* plays an indispensable role in the onset of embryonic stem cells (ESCs) differentiation [26]. SLC35B4 (solute carrier family 35, member B4) regulates obesity and glucose homeostasis [27].

#### 3. Density peaks located the ICR/gDMR of Zac1 and candidate ICRs for several potential imprinted genes

The ICR of *Zac1* is in Chr10qA. This chromosomal band covers 4 subbands and contains ∼ 34 Mb DNA. For that DNA section, we observed 7 robust peaks (1 per ∼ 4.8 Mb). The *Plagl1* locus includes an intragenic conserved CGI near one of the *Plagl1* transcripts known as *Zac1*. In oocytes, this CGI is methylated and regulates parent-of-origin specific gene expression [28]. A previous study found a large cluster of ZFBS-Morph overlaps in the *Zac1* ICR/gDMR [13]. The density-plot revealed a very robust density peak that encompasses that cluster in the intragenic CGI that corresponds to the *Zac1* ICR/gDMR (Fig. S5). In subband 10qA1, a candidate ICR includes the promoter and the TSS of the longest *Zbtb2* transcript (Fig. S6). ZBTB2 (Zinc Finger and BTB domain containing 2) is a transcription factor that is ‘a reader’ of unmethylated CpG island promoters to regulate ES cells differentiation in mice [29]. Subband 10qA3 covers an intergenic candidate ICR between the TSSs of *Nhsl1* and *Ccdc28a* (Fig. S7). According to GENE at NCBI, *Ccdc28a* (coiled-coil domain containing 28A) is expressed in adult testis (https://www.ncbi.nlm.nih.gov/gene/215814#gene-expression). The function *Nhsl1* (NHS like 1) is not clearly understood. Subband 10qA4 includes a candidate ICR in a CGI that encompasses the TSSs of *Stx7* transcripts (Fig. S8). Syntaxin-7 is among a group of proteins that function in vesicle transport and fusion events [30].

Besides the known ICR/gDMR of *Zac1*, subband 10qA2 includes 3 candidate ICRs (Fig. 4). They map to *Cited2* (Fig. S9), to a CGI associated with several *Hivep2* transcripts (Fig. S10), and to an uncharacterized gene (*AK045519*). *Hivep2* (human immunodeficiency virus type I enhancer-binding protein 2) encodes a transcription factor that binds DNA [31]. In mice, *Hivep2* deficiency Induced mild chronic inflammation in the brain and conferred molecular, neuronal, and behavioral phenotypes related to schizophrenia [32]. *Cited2* (CBP/p300-interacting transactivator 2) also is involved in the regulation of transcription. Absence of *Cited2* in mouse embryos caused congenital heart disease by perturbing left-right patterning of the body axis [33].

**Figure 4.**
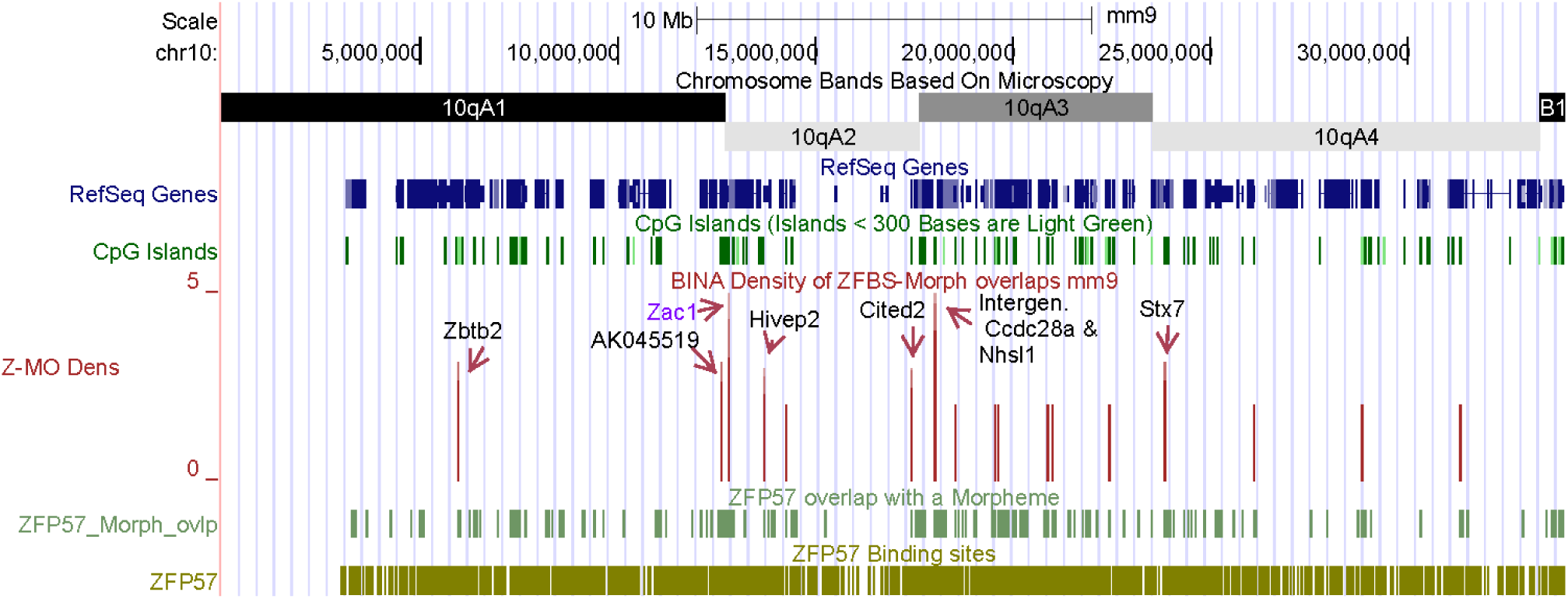
A long DNA section encompassing several candidate ICRs and the known ICR/gDMR of *Zac1*. A robust density peak pinpointed the known ICR/gDMR of *Zac1* in ∼ 34 Mb long DNA. Several additional robust peaks mark the positions of candidate ICRs for potential imprinted genes.

#### 4. Density peaks revealed the known ICR/gDMR of Igf2r – Airn imprinted domain, and candidate ICRs mapping to Arid1b and to the tctex-1 locus (a candidate gene family for a mouse t complex sterility)

In evaluations, we closely inspected the density-plot for every mouse autosomal chromosome. For example, examine the positions of peaks in the plot that covers the entire Chr17 (Fig. 5). Even along the entire chromosomal DNA (∼ 95.3 Mb), we could detect the robust density peak that corresponded to ICR of the *Igf2r* – *Airn* imprinted domain. Located in the 2^nd^ intron of *Igf2r*, the ICR/gDMR of the domain regulates parent-of-origin specific expression of several genes [34]. This intronic ICR/gDMR is in a CGI in the vicinity of *Airn* promoter [16]. Also known as *Air, Airn* is the only paternally expressed gene in that imprinted domain. Transcription of *Airn* produces a noncoding RNA that is antisense orientation with respect to *Igf2r* and *Mas* [35]. In the density-plots, a very robust density peak maps to the intragenic CGI that regulate the expression of genes in the imprinted domain (Fig. S11). This observation supports the hypothesis that robust density peaks could correspond to the ICRs in the mouse genome.

**Figure 5.**
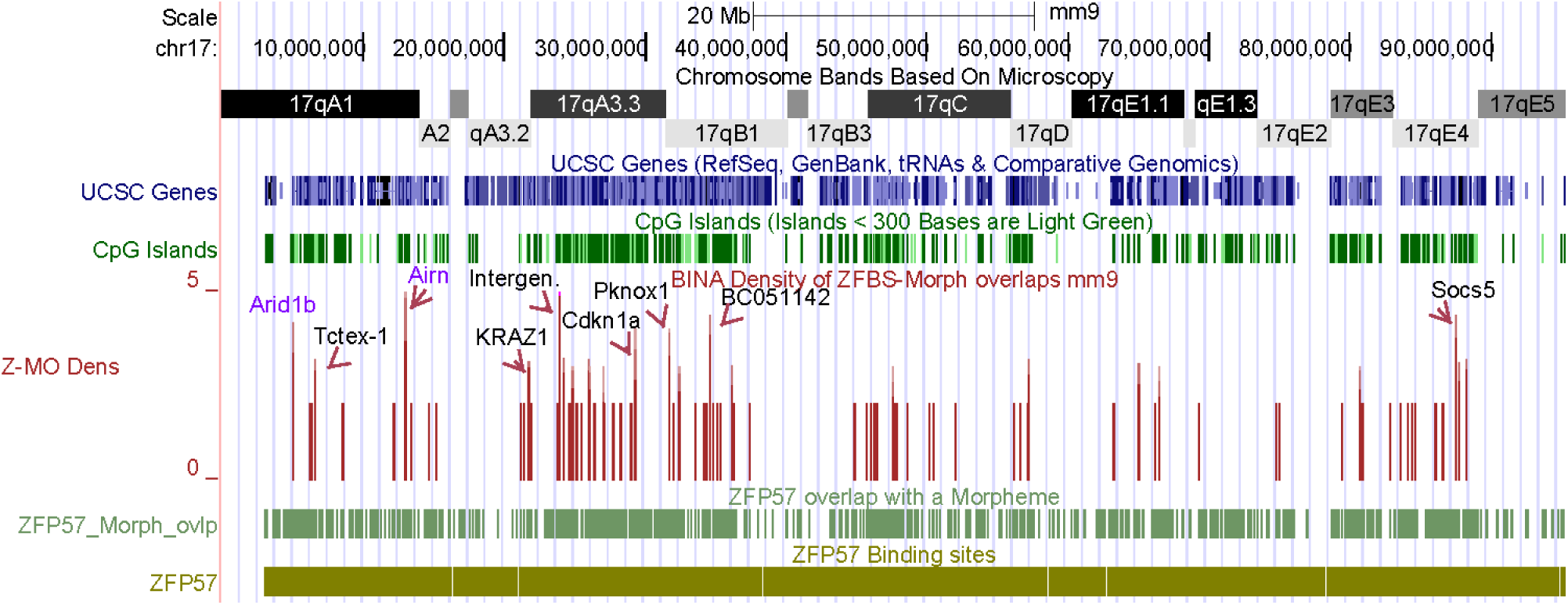
The position of peaks in the density-plot of Chr17. By uploading our data onto the UCSC genome browser, we could view the positions of density peaks across an entire chromosome. Peak-heights reflect the number of ZFBS-morph overlaps within the sliding 850-base window. A large fraction of peaks corresponds to those covering 2 ZFBS-morph overlaps. Robust peaks are less frequent and distributed unevenly along the DNA. In many regions, they appear as clusters –possibly a reflection of the intricacies of epigenetic regulation in clusters [36]. In exploratory evaluations, we encountered instances where 2 closely spaced peaks produced a robust density peak.

Key features of the genome browser include the option of uploading files to create custom tracks, as shown in this report, and to enhance the resolution of peaks by zooming in to obtain enlarged or close up views [37-39]. For example, an enlarged view shows the position of density peaks in a DNA section that covers the subbands in Chr17qA (Fig. 6). The displayed view covers ∼ 23 Mb DNA and 4 robust density peaks (1 / 5.75 Mb). One peak corresponds to the known ICR/gDMR of the *Igf2r* – *Airn* imprinted domain. Three peaks mark the positions of candidate ICRs. They are in the vicinity of *Arid1b, KRAZ1*, and *tctex-1* loci (Fig. 6). We chose that section because it includes the ICR/gDMR of the *Igf2r* – *Airn* imprinted domain and a candidate ICR for *Arid1b* –a known imprinted gene in mouse [40]. A robust density peak maps to the longest *Arid1b* transcripts (Fig. 7). *Arid1b* (AT rich interactive domain 1B) encodes an enzyme (KDM5B) that removes methyl-marks from H3K4. Depletion of *Arid1b* in ES cells caused defects in gene expression programs [41].

**Figure 6.**
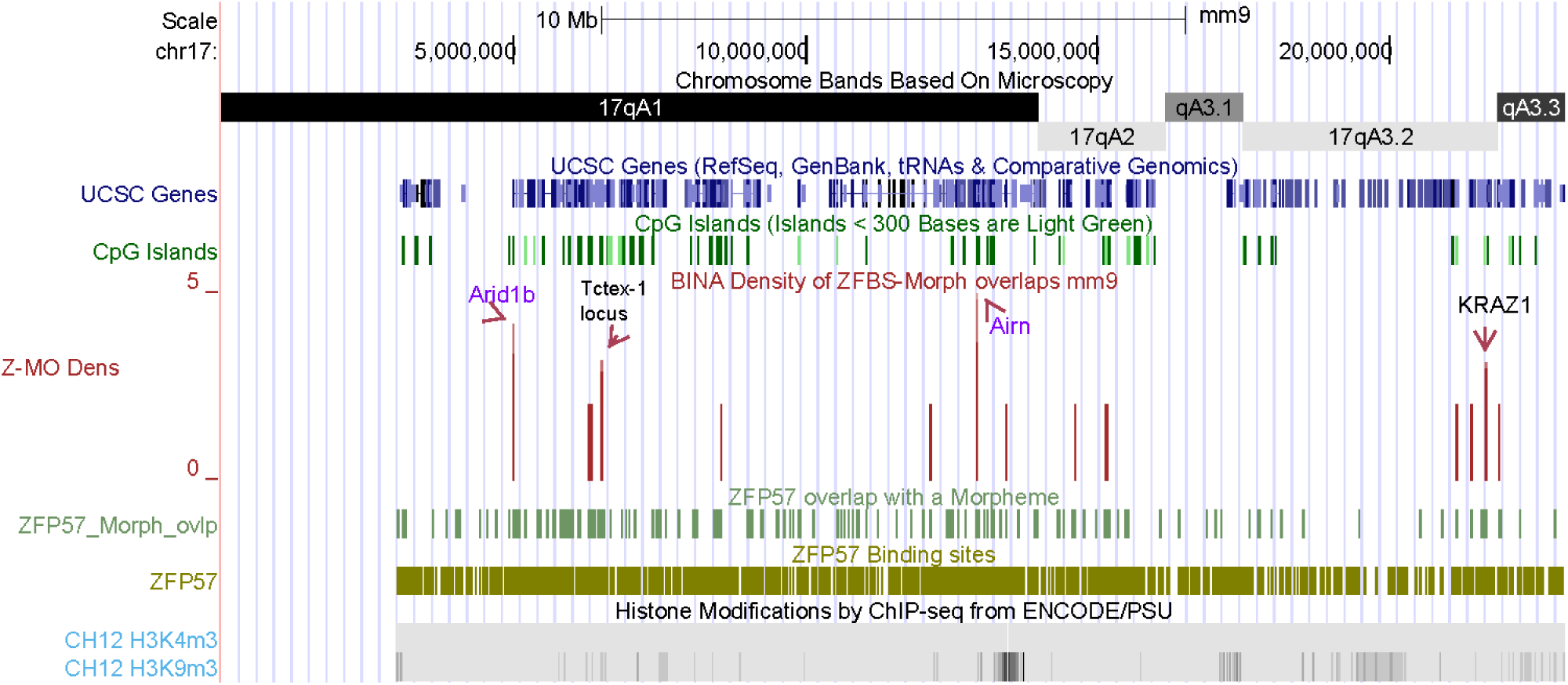
A view of the positions of the robust density peaks in the proximal end of Chr17. This chromosomal section includes candidate ICRs for *Arid1b* and potential imprinted loci.

**Figure 7.**
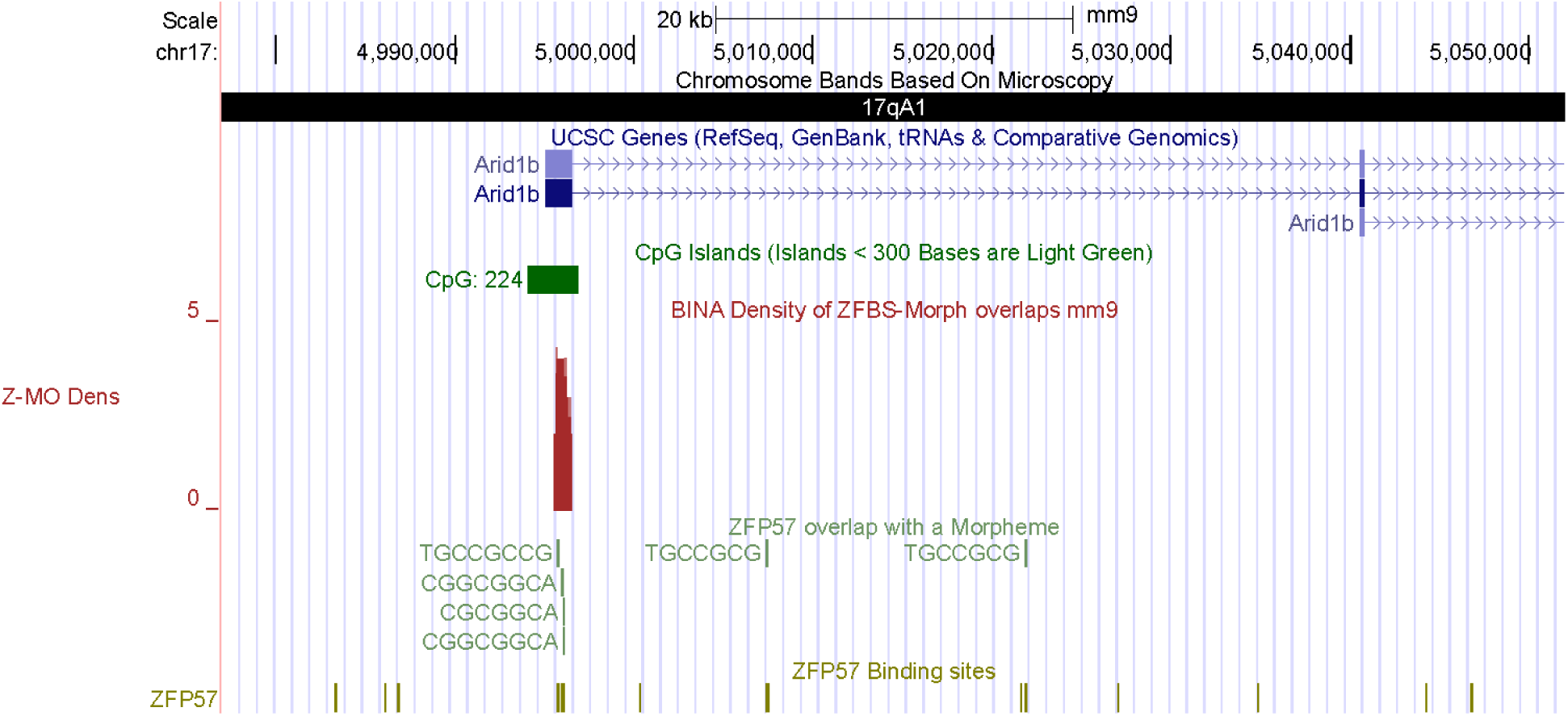
The position of a candidate ICR for *Arid1b* –a known imprinted gene in mouse [40].

In Chr17qA, another robust density peak is in the vicinity of *KRAZ1* (Figs. 5, 6, and S12). *KRAZ1* is in a chromosomal section that encompasses several genes for proteins whose structure includes zinc fingers (Fig. S12). *KRAZ1* encodes a protein with Krüppel-type zinc finger structural motif also found in ZNF57, KRAZ2, and ZNF445; references [42-44]. As ZNF57, both KRAZ1 and KRAZ2 interacts with KAP1 [43]. Also known asTRIM28, KAP1 plays a central role in silencing of the imprinted genes [6].

Notably, Chr17qA is in the chromosomal section that includes naturally occurring variants known as t haplotypes [45]. The variants in t haplotypes are alleles of genes that exist as a polymorphism in wild mouse populations [46]. While males heterozygous for a t haplotype are normal, homozygous males are sterile [46, 47]. All t haplotypes contain rearranged segments of DNA known as the t complex [45]. The aberrant expression of *tctex-1* is solely dependent on the t haplotype genes and occurs only in germ cells [45]. The *tctex-1* locus consists of a four member multigene [48]. In males, the wild-type *tctex-1* transcript was first abundantly expressed at the pachytene stage of meiosis and persisted throughout spermatogenesis [48]. In the density plot of Chr17, a robust density peak is in the vicinity of *tctex-1* (Figs. 5 and 6). Members of this gene family includes *Dynlt1* (dynein light chain Tctex type 1). Dyneins are large multicomponent microtubule-based molecular motors involved in many fundamental cellular processes including mitosis [49]. In the *tctex-1* locus, a candidate ICR maps to a CpG island at ∼ 800 bps upstream of the TSS of one of the *Dynlt1e* genes/transcripts (marked by an arrow in Fig. 8). Thus, our finding predicts that the *tctex-1* locus could be an imprinted domain regulated by an ICR defined by a robust density peak (Figs 5, 6, and 8).

**Figure 8.**
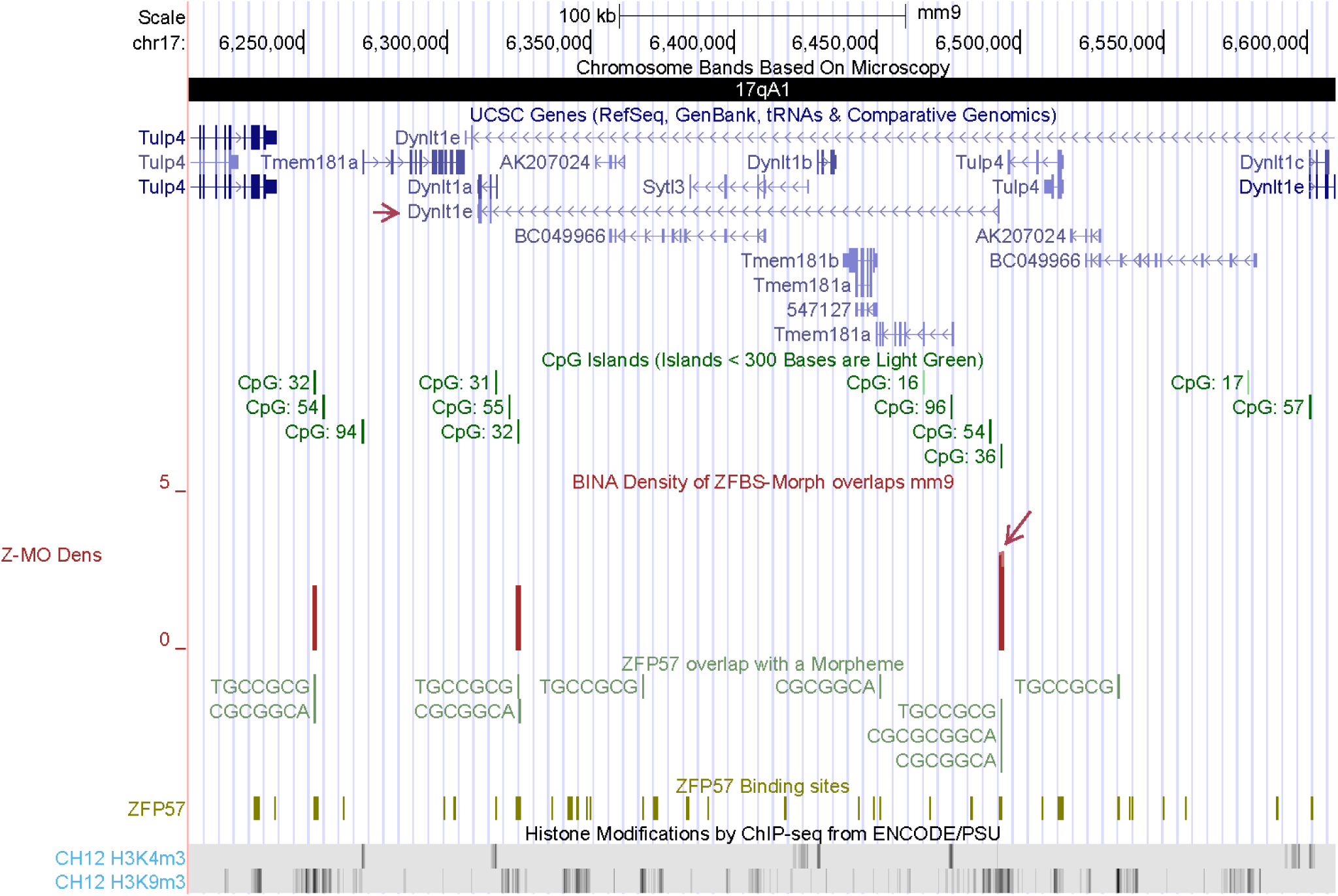
A candidate ICR mapping to the *tctex-1* locus. An arrow points to the robust density peak providing a candidate ICR for regulation of expression from the *tctex-1* locus. This peak is upstream of TSS of one the genes designated as *Dynlt1e* (marked by an arrow). Noteworthy are several patches of repressive H3K9me3 marks dispersed across the locus. These marks reinforce our prediction that the *tctex-1* locus could be an imprinted domain.

#### 5. Density peaks revealed the known ICR/gDMR of Nnat, and candidate ICRs for Bcl2l1 and two potential imprinted genes

In our exploratory analyses, we also examined an ∼ 12.6 Mb DNA section encompassing Chr2qG3 and Chr2qH1 (Fig. 9). In that DNA section, we observed 4 robust density peaks (1 peak / 3.15 Mb). One peak pinpointed the ICR of *Nnat*. This ICR is in one of the introns of gene of a gene (*Blcap*) that is biallelically expressed [50, 51]. With respect to *Blcap, Nnat* is transcribed in the opposite direction. In transgenic animals, the ICR/gDMR of *Nnat* was mapped to a CpG island spanning the 1^st^ exon and extending into intron 2 [52]. According to genomic maps, from *Nnat* are produced several transcripts with differential exon utilization patterns (Fig. S13). In close-up views, we observed 2 peaks within a DNA segment that encompasses the intragenic *Nnat* ICR (Fig. S13). Besides the peak corresponding to the known, the ∼ 12.6 Mb DNA includes 3 additional robust peaks. Two peaks are in Chr2qH1. They are in the vicinity of *Bcl2l1* and *Ccm2l*. The 3^rd^ peak maps to Chr2qG3, in the vicinity of *Nkx2-4* (Fig. 9). In critical evaluations of imprinted gene expression by RNA–Seq, *Bcl2l1* was listed as a candidate imprinted gene [53]. Furthermore, evidence supports the unequal contribution of the *Bcl2l1* paternal and maternal alleles to brain development. Also referred to as *Bcl-x*, haploinsufficiency of *Bcl2l1* led to male-specific defects in fetal germ cells [54]. Notably, expression was paternally-biased for the longest *Bcl2l1* isoform [55]. These findings support our prediction of a candidate ICR in the CGI that is associated with the longest *Bcl2l1* transcript (Fig. 10). Furthermore, the predicted ICR is within chromatin boundaries consisting of CTCF, RAD21, and SMC3.

**Figure 9.**
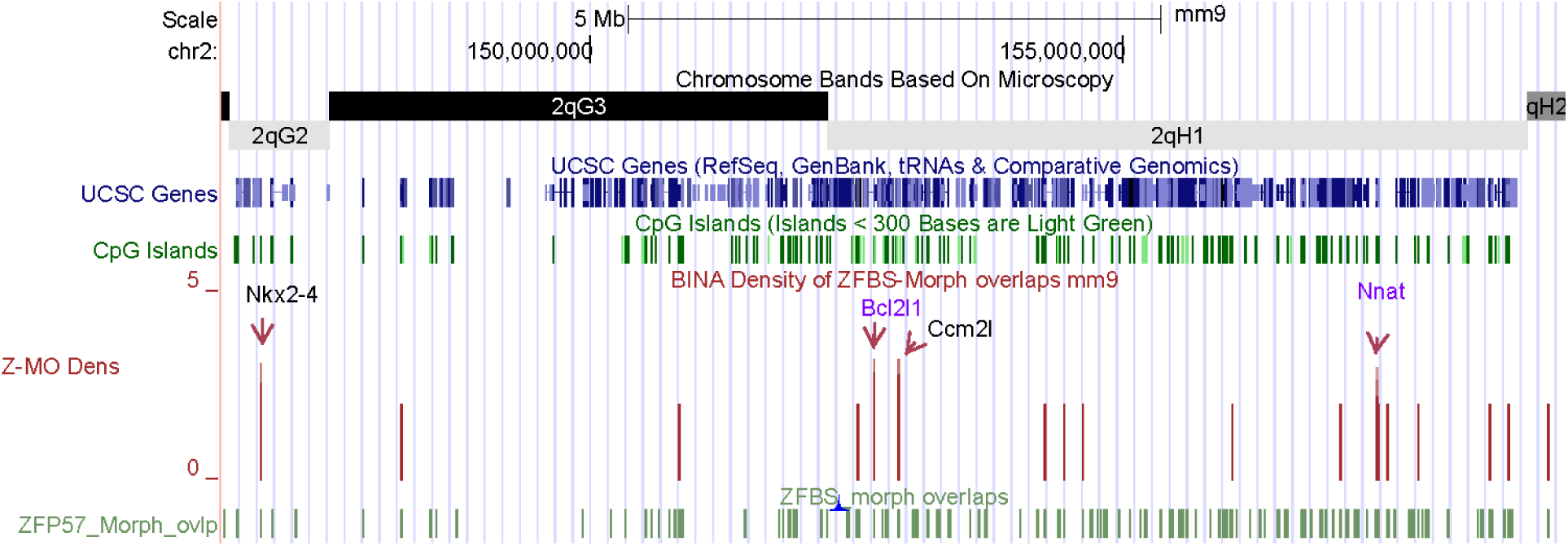
Genomic positions of density peaks along Chr2qG3 and Chr2qH1. An arrow points to the peak that corresponds to the known ICR/gDMR of *Nnat*. Additional arrows mark the position of candidate ICRs.

**Figure 10.**
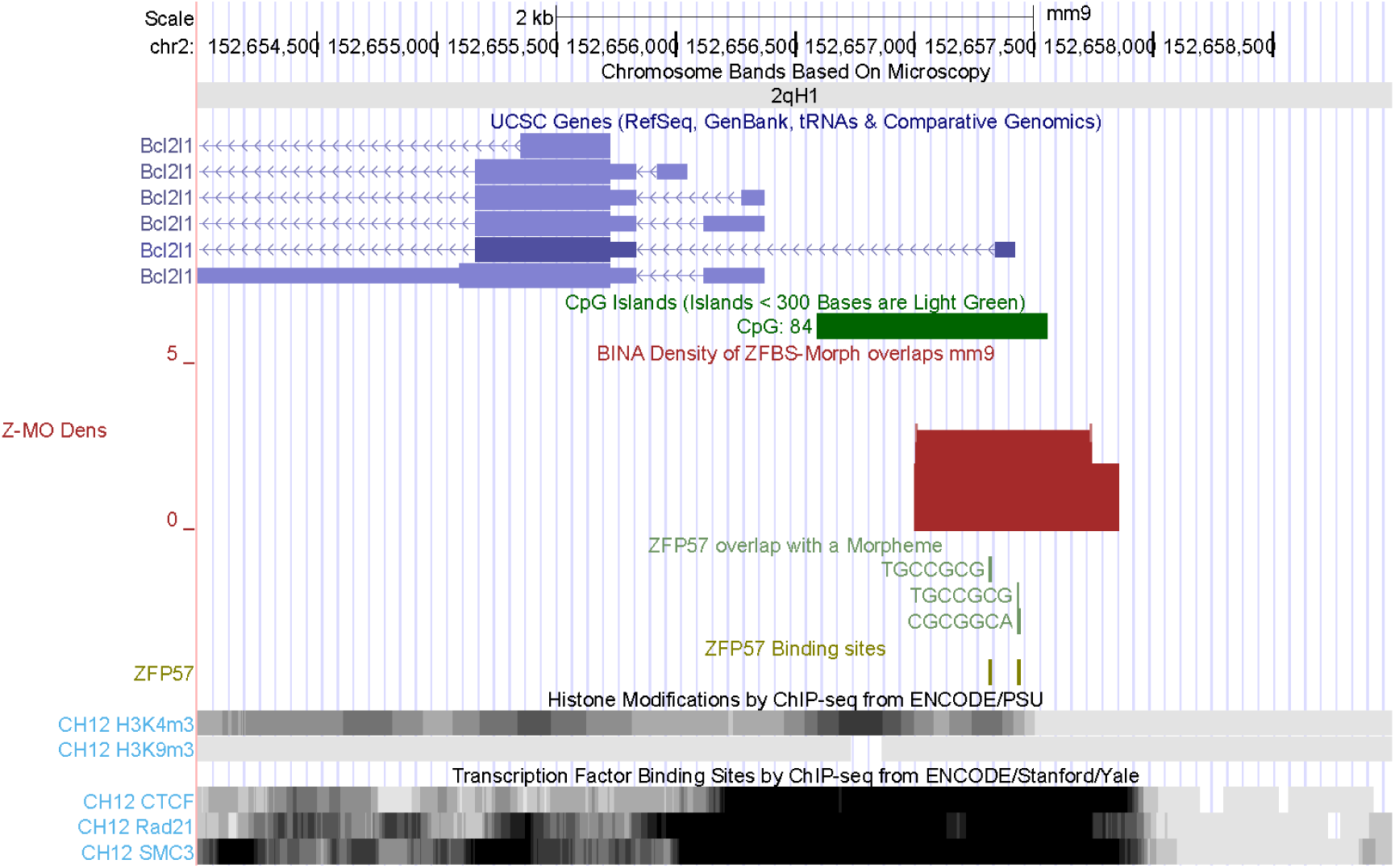
The position of a candidate ICR with respect to the longest *Bcl2l1* transcript. Expression analyses have identified *Bcl2l1* as a candidate imprinted gene [53].

The candidate ICR for *Ccm2l* is intragenic and corresponds to a density peak is in the 5^th^ exon of the gene (Fig. S14). *Ccm2l* (cerebral cavernous malformation 2-like) is a paralog of *Ccm2* [56]. CCM2 functions in the CCM signaling [57]. Its paralog (CCM2L) is selectively produced in endothelial cells during angiogenesis [57]. *Ccm2l* ^-/-^ animals exhibited embryonic lethality at E11 associated with myocardial thinning [56]. In Chr2qG3, a robust density peak maps to *Nkx2-4* (Fig. 9). This candidate ICR is in a CpG island that encompasses *Nkx2-4* TSS (Fig. S15). Notably, a chromatin section within *Nkx2-4* includes repressive H3K9m3 marks. *Nkx2-4* (NK2 homeobox 4) encodes a transcription factor whose structure includes a homeodomain for binding DNA [58]. During mouse embryogenesis, *Nkx-2.2* transcripts were dispersed in localized domains possibly for specifying diencephalic neuromeric boundaries [58].

### Several candidate ICRs are in the vicinity of chromatin boundaries

It is well known that the ICR of the *H19 – Igf2* imprinted domain includes chromatin boundaries [20, 21]. Furthermore, several of the known ICRs regulate a subset of gene-transcripts; *i.e., Zac1, Inpp5f_v2*, and *Mest* [28, 59, 60]. Therefore, to further evaluate our approach, we inspected the density-plots to determine whether any of the candidate ICRs mapped to specific gene-transcripts, to chromatin boundaries, or to both. For these studies, we surveyed the density-plots with respect to results of ChIPs obtained for CTCF, RAD21, and SMC3 – reference [61]. Within the sampled DNA sections, we found several instances where a candidate ICR was associated with a single transcript and a chromatin boundary. Examples for a single transcript include a candidate ICR in a segment encompassing several members of the *Six* family (Fig. S16). The corresponding density peak is in a CGI that encompasses the TSS and the 1^st^ exon of *Six1*. It maps to a region that includes chromatin boundaries consisting of CTCF, RAD21, and SMC3 (Fig. S16). *Six1* (sine oculis-related homeobox 1) encodes a transcription factor that plays central roles in the pre-placodal ectoderm, a common field from which arises all cranial sensory placodes including adenohypophyseal, olfactory, lens, trigeminal, epibranchial, and otic [62].

Examples of candidate ICRs mapping to a subset of gene transcripts include a density peak near the 5’ end of the longest *Hectd1* transcriptional variants (Fig. S17). This peak maps to a CGI that encompasses chromatin boundaries. *Hectd1* (HECT domain E3 ubiquitin protein ligase 1) is required for development of the junctional zone of the placenta. Disruption of *Hectd1* resulted in mid-gestation lethality and intrauterine growth restriction [63]. Another example is the candidate ICR that maps to the longest *Bcl2l1* transcript (Fig. 10). Functionally, BCL2L1 belongs to a group of proteins related to BCL-2 (B cell leukemia/lymphoma 2). Members of this group control cell death through complex interactions that regulate mitochondrial outer membrane permeabilization. This process leads to the irreversible release of intermembrane space proteins with subsequent caspase activation and apoptosis [64].

### Several candidate ICR map to the vicinity of genes encoding transcription factors and proteins involved in sex-related processes

Since transcription factors regulate the expression of many genes, they play central roles in numerous physiological processes. In humans, mutations in transcription factor genes could cause developmental abnormities know as syndromes [65]. As summarized above, near several of the candidate ICRs, we found genes encoding regulators of transcription (*i.e. Taf4a, Zbtb2, Six1, Hivep2, Nkx2-4*). Additional examples include *Nfix, Maml3, Pou3f3, Foxo6* and *Pou3f1. Nfix* is in Chr8qC3. Density-plots revealed a candidate ICR near the two longest *Nfix* transcripts (Fig. S18). *Nfix* (nuclear factor I/X) belongs to a group of transcription factors that play pivotal role in the development of the nervous system [66]. *Maml3* (mastermind like transcriptional coactivator 3) specifies a transcriptional coactivator. A candidate ICR is located in a CGI that encompasses the TSS of the longest *Maml3* transcript (Fig. S19). *In vivo, Maml3* and one of its paralogs (*Maml1*) are essential for Notch signaling. This system plays a pivotal role in metazoan development [67]. Even though *Maml3*-null mice showed no apparent abnormalities, mice null for both *Maml1* and *Maml3* died early in the organogenic period and displayed classic pan-Notch defects [67]. A relatively long DNA section (∼ 20 Mb) that encompasses Chr1qB includes only 2 robust density peaks (1 / 10 Mb). These 2 candidate ICRs are in the vicinity of *Pou3f3* and *Rev1* (Fig. 12). One is in a CGI that encompasses *Pou3f3* TSS (Fig. S20). The displayed segment includes several repressive H3K9me3 marks. The other is in a CGI that encompasses *Rev1* TSS (Fig. S21). This CGI is in the vicinity of chromatin boundaries. POU3F3 or BRN1 belongs to a group of transcription factors that influence neurogenesis, molecular identity, and migratory destination of upper-layer cells of the cerebral cortex [68]. Their structure incudes a POU-homeodomain for binding DNA [69].

**Figure 11.**
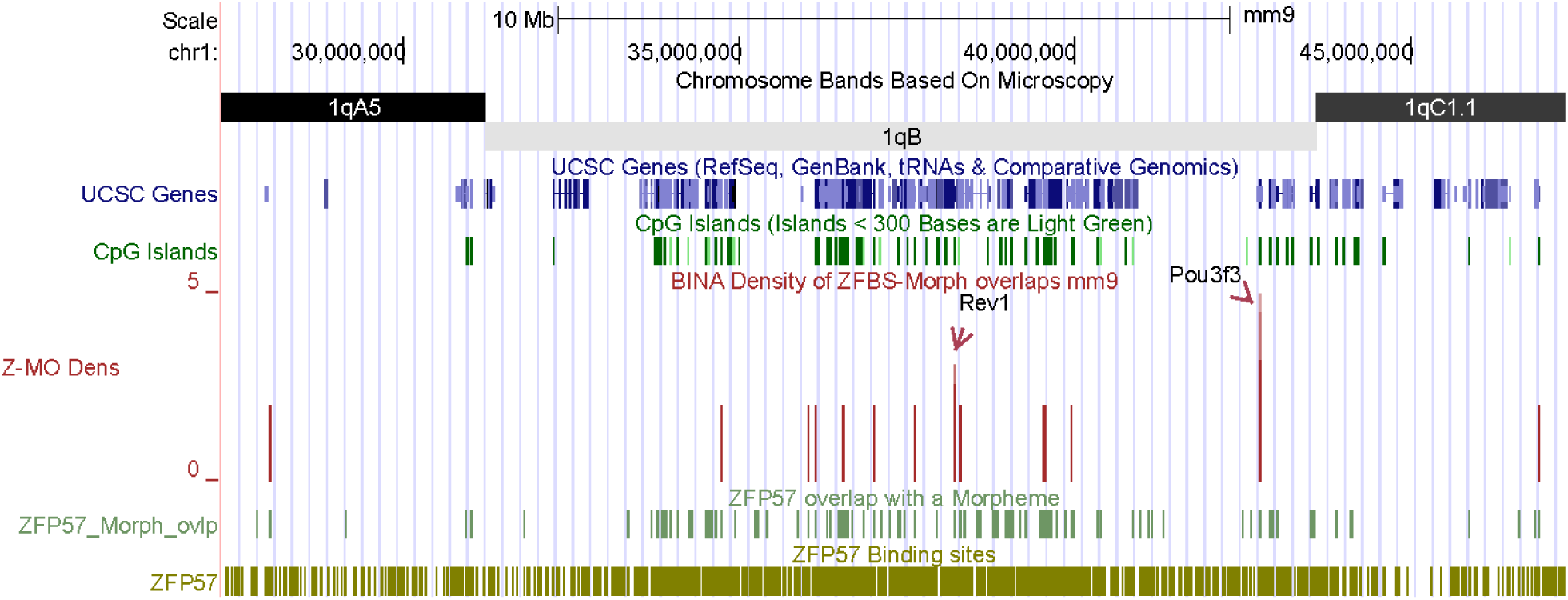
Two candidate ICRs within a relatively long DNA section. They are in the vicinity of *Rev1* and *Pou3f3*.

**Figure 12.**
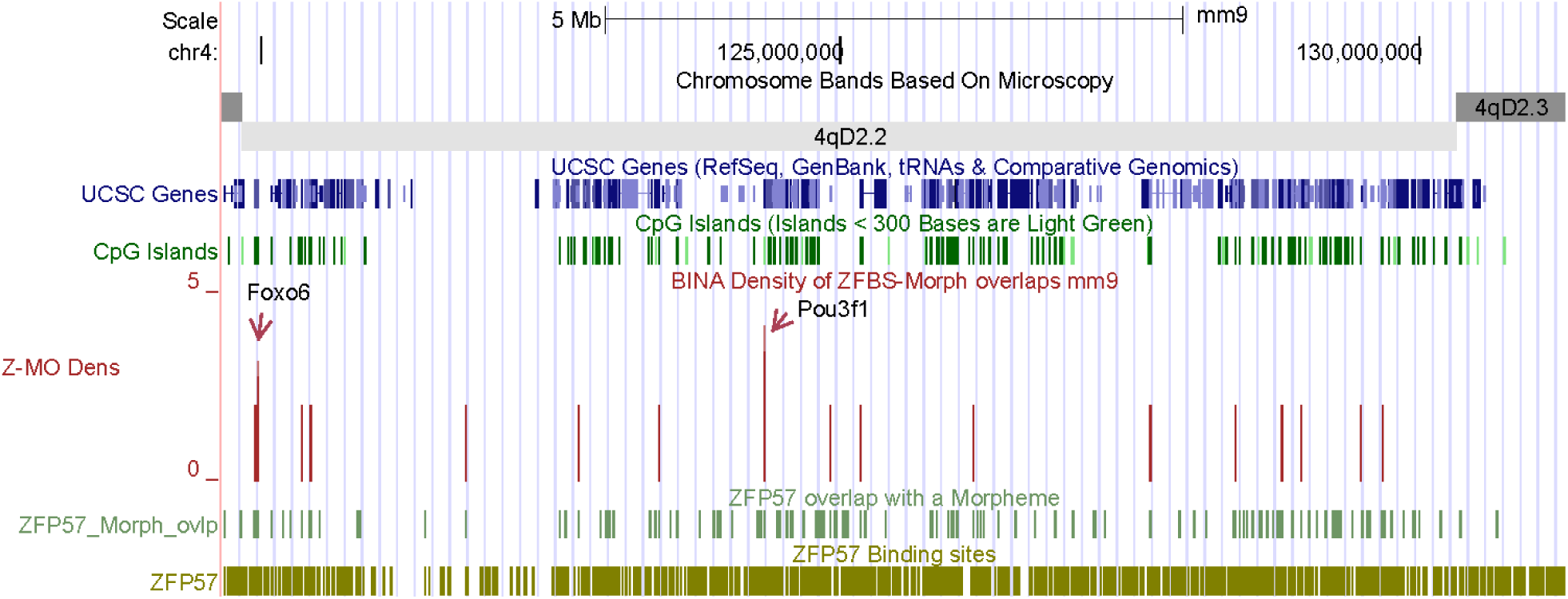
Two candidate ICRs mapping to two transcription factor genes (*Foxo6* and *Pou3f1*).

*Foxo6* and *Pou3f1* are in Chr4qD2.2 (Fig. 12). The displayed DNA section (∼11.6 Mb) encompasses 2 robust density peaks (1 peak per 5.8 Mb). One peak gives a candidate ICR in a CGI that contains Foxo*6* TSS and promotor (Fig. S22). The other corresponds to a candidate ICR in a CGI that encompasses *Pou3f1* TSS (Fig. S21). The structure of FOXO6 includes a forkhead (winged helix) domain for binding *cis* elements in DNA [70]. That of POU3F1 includes a POU-homeodomain for interactions with DNA [69]. Even though both FOXO6 and POU3F1 are transcription factors, they play distinguishable roles in the regulation of gene expression. *Foxo6* is expressed at late stages of face development [71]. Foxo6 ^-/-^ mice underwent expansion of the face, frontal cortex, olfactory component and skull [71]. *Pou3f1* plays a role in spermatogenesis [72]. Briefly, fertility of adult males requires self-renewal and differentiation of spermatogonial stem cells. A study found that in cells cultured *in vitro, Pou3f1* expression was up-regulated in response to extrinsic stimulation with GDNF (glial cell line-derived neurotrophic factor). In cross-sections of prepubertal and adult testes, biochemical-analyses localized POU3F1 to spermatogonia. A reduction in *Pou3f1* expression induced apoptosis of cultured germ cells [72].

In fact, in our studies we came across more than a few candidate ICRs that mapped to the vicinity of transcripts involved in sex-related processes in males and females. In previous sections we gave a few examples. Additional examples include *Mgea5, Cnnm1, Chd5*, and *BC051142* (Figs. S24-S27). Also known as *OGA*, MGEA5 influences folliculogenesis [73]. Expression of *Cnnm1* (cyclin M1) is associated with cell cycle and differentiation in spermatogenic cells in mouse testis [74]. *Chd5* expression is restricted to post-meiotic spermatids [75]. CHD5 (chromodomain helicase DNA binding protein 5) mediates histone-to-protamine replacement impacting the cascade of molecular events underlying chromatin remodeling during spermatogenesis [75]. *Chd5* expression peaks just as the most dramatic chromatin remodeling starts to take place [76]. *BC051142* function is unknown. It is among the 54 evolutionarily-conserved and testis-enriched genes uncovered by genome engineering techniques [77]. Furthermore, according to GENE at NCBI, *BC051142* expression is restricted toward adult mouse testes (https://www.ncbi.nlm.nih.gov/gene/?term=BC051142#gene-expression).

## DISCUSSION

To fully understand the developmental consequences of uniparental embryos, it is necessary to obtain a complete list of all imprinted genes [78]. Towards this goal, we developed a bioinformatics strategy. We built our approach on results showing that ∼ 90% of the known ICRs/gDMRs included clusters of ZFBS-morph overlaps [13, 16]. In order to remove background noise, we created plots to display the density of ZFBS-morph overlaps along genomic DNA. Previously, a study assessed the predictive potential of this strategy by examining the density-plots obtained for the human genome [19]. In this report, we assessed whether this approach also could be applied to the discovery of candidate ICRs and imprinted genes in mouse. To explore that question, initially we investigated whether we could locate known ICRs/gDMRs in relatively long DNA sections. We found that in plots, robust density peaks clearly pinpointed the essential ICR of the *Gnas* complex locus in 8.25 Mb DNA (Fig. 2), the intragenic ICR of *Mest* in 36 Mb DNA (Fig. 3), the ICR of *Zac1* in 34 Mb DNA (Fig. 4), the ICR of the *Igf2r* – *Airn* imprinted domain in 23 Mb DNA (Fig. 5), and the ICR of *Nnat* in 12.6 Mb DNA (Fig. 6). Based on the above and related observations, we deduced that in the density-plots, robust peaks reflected the position of actual or candidate ICRs. We posited that the genes in the vicinity of candidate ICRs corresponded to potential imprinted genes. To explore these ideas, we sampled several of the robust peaks to inspect their positions with respect to genomic landmarks, including genes, transcripts, and the CpG islands.

### 1. The majority of the candidate ICRs maps to CGIs; a subset corresponds to specific gene-transcripts; a smaller subset corresponds to regions associated with proteins that produce chromatin boundaries

Many of the known ICRs coincide with CGIs [5]. Similarly, we found that in the density-plots, the analyzed robust peaks primarily mapped to CGIs. These islands are dispersed in various genomic locations including 1^st^ exons, near TSSs, and within intra and intergenic sequences. In many cases, a known ICR/gDMR represses the expression of one transcript or a group of transcripts [60, 79, 80]. Similarly, we found candidate ICRs that mapped to a single of a subset of gene-transcripts. Examples for a single transcript includes *Arid1b, Nkx2-4, Six*1, and *Pou3f1* (Figs. 7, S15, S16 and S21). Examples for a subset of transcripts include the predicted ICRs that map to *Bcl2l1, Taf4a, Lsm14b, Impdh1, Zbtb2*, and *Nfix* loci (Figs. 10, S1, S2, S6, S18). In a few cases, a known ICR/gDMR is intergenic. Similarly, we found candidate ICRs within intergenic sequences (*i.e*., Figs. S4 and S7). Also, in a few instances, a known ICR/gDMR contains sites for binding CTCF to produce chromatin boundaries [20, 21]. Through association with CTCF, Rad21 and SMC3 contribute to the establishment or maintenance of topological domains [81]. In our analyses we found a few candidate ICRs that mapped to or near chromatin boundaries consisting of CTCF, with or without Rad21 and SMC3. Examples include *Bcl2l1, Impdh1, Cited2, Six1*, and *Hectd1* (Figs. 10, S2, S9, S16, and S17).

### 2. Several of the candidate ICRs map to genes with functions ranging from developmental to sex-related processes

Many of the imprinted genes are involved in regulation of embryonic growth and developmental processes. Others play key roles in neurological processes and behavior [82]. Therefore, for our studies we conducted literature surveys to investigate the function of the genes in the vicinity of the candidate ICRs that we examined. In the context of embryonic stem cells (ESCs), we obtained *Zbtb2, Fam40b/Strip2*, and *Arid1b*. In cells, ZBTB2 interacted with CpG island promoters, where it acted as a transcriptional activator [29]. STRIP2 was required for lineage commitment of murine ESCs [26]. ARID1B affected gene expression by catalyzing the removal of H3K4methyl marks from chromatin [83]. In the context of fetal growth, we obtained *Hectd1*. This gene encodes an enzyme required for the development of the junctional zone of the placenta. Disruption of *Hectd1* resulted in mid-gestation lethality and intrauterine growth restriction [63].

In the context of developmental processes, we obtained several genes encoding transcription factors – *i.e*., *Cited2* and *Foxo6. Cited2* impacts left-right patterning of the body’ s axis [33]. Functions of FOXO6 include regulation of Hippo signaling and the growth of the craniofacial complex [71]. In the context of organogenesis, we obtained *Maml3*. Even though *Maml3*-null mice showed no apparent abnormalities, mice null for both *Maml1* and *Maml3* died early in the organogenic period [67]. In the context of vision, we obtained *Impdh1*. Mutations in the corresponding human gene caused retinitis pigmentosa –an inherited retinal degeneration characterized by the early onset of night blindness followed by a progressive loss of the visual field [24, 25]. In the context of sensory organs, we obtained *Six1*. This gene specifies a transcription factor that plays central roles in the pre-placodal ectoderm [62]. In the context of the brain and neuronal functions, we obtained *Nfix, Pou3f3*, and *Hivep2*. NFIX contributes to the development of the neocortex, hippocampus, and cerebellum [84]. POU3F3 influences neurogenesis, molecular identity, and the migratory destination of upper-layer cells of the cerebral cortex [68]. *Hivep2* deficiency confers molecular, neuronal, and behavioral phenotypes related to schizophrenia [32].

In the context of sex-related processes, we obtained several genes. *Lsm14b* and *Mgea5* impact oogenesis. *Lsm14b* encodes an RNA binding protein with an essential role in oocyte meiotic maturation [23]. Also known as OGA, MGEA5 catalyzes the removal of N-acetylglucosamine from serine and threonine residues in many proteins [85]. Transcriptome profiling identified this enzyme as an upstream regulator of granulosa cells of bovine ovarian follicles [73]. For spermatogenesis, we obtained *Pou3f1, Cnnm1, Chd5*, an uncharacterized gene (*BC051142*), and *tctex-1* locus. POU3F1 is involved in survival and self-renewal of mouse spermatogonial stem cells [72]. CNNM1 is produced in mouse testes from neonatal to adult stages [74]. CHD5 mediates histone-to-protamine replacement and impacts the cascade of molecular events underlying chromatin remodeling during spermatogenesis [75]. Among sex-related loci, the most intriguing could be the *tctex-1* locus within the proximal half of Chr17 [48]. The t-haplotypes include this chromosomal section –They are variant alleles of genes linked together by four inversions [46, 47]. While females carrying two t-haplotypes are fertile, males are sterile. The spermatozoa in males exhibit motility defects, rendering them unable to penetrate zona pellucida-free oocytes [46, 47]. *Dynlt1e* is among the genes in *tctex-1* –a candidate gene family for a mouse t complex sterility [48]. Our data indicate that in the proximal half of chromosome 17, the *tctex-1* locus might be regulated by genomic imprinting through a candidate ICR that maps to a CGI upstream of one of the *Dynlt1e* genes/transcripts. Consistent with this interpretation are several H3K9me3 marks dispersed across the entire *tctex-1* locus (Fig. 8).

## 4. CONCLUSION

This report offered a strategy for simultaneous discovery of candidate ICRs and imprinted genes in the mouse genome. With this approach, we could clearly discern the genomic positions of several of the known ICRs/gDMRs within relatively long DNA sections. Furthermore, with respect to predicted ICRs, we could locate potential imprinted genes/transcripts with diverse and physiologically important functions –including fetal grow, organogenesis, craniofacial feature, patterning of the body axis, development of sensory organs, several sex related processes, and chromatin remodeling. These and related findings could lend further support for robustness of our strategy. Nonetheless, only experimental validations could demonstrate the strength of our approach. Therefore, we offer links for accessing and downloading our data on the positions of ZFBS and ZFBS-Morph overlaps [86], peaks in the density-plots [87], and the MLL1 morphemes in the build mm9 of the mouse genome [88]. Also available are links for downloading data-files obtained for the build hg19 of the human genome [89-91].

## 5. METHODS

Previously, two reports gave links for accessing the genomic positions of a set of composite DNA elements consisting of the ZFP57 binding site overlapping a subset of the MLL1 morphemes [13, 86]. We wrote a Perl script for creating the density-plots. The script opened the file containing the positions of ZFBS-Morph overlaps in a specified chromosome. Next, the script scanned the file to count and to report the number of ZFBS-Morph overlaps within a sliding window consisting of 850 bases. By omitting isolated occurrences, the script removed background noise. Subsequently, we combined and tailored the outputs of the program for upload as a custom track at the UCSC genome browser. Several publications offer concise overviews about how to use the UCSC browser [38, 39]. Default tracks at the browser include the positions of genes, transcripts, the CpG islands, SNPs, and the ENCODE data [37, 61]. The positions of SNPs could facilitate designing primers to test the validity of our predictions. With the ENCODE data, one could examine the candidate ICRs in the context of histone marks, DNase I hypersensitive sites, and results of ChIPs determined for several nuclear proteins including CTCF, RAD21, and SMC3; for details, see reference [92].

## Competing interests

none

## Acknowledgments

We thank Zena Narod Lamp for editing our manuscript, Arnold Stein for helpful discussions.

## SUPPLEMENTAL FIGURES

**Fig. S1.**
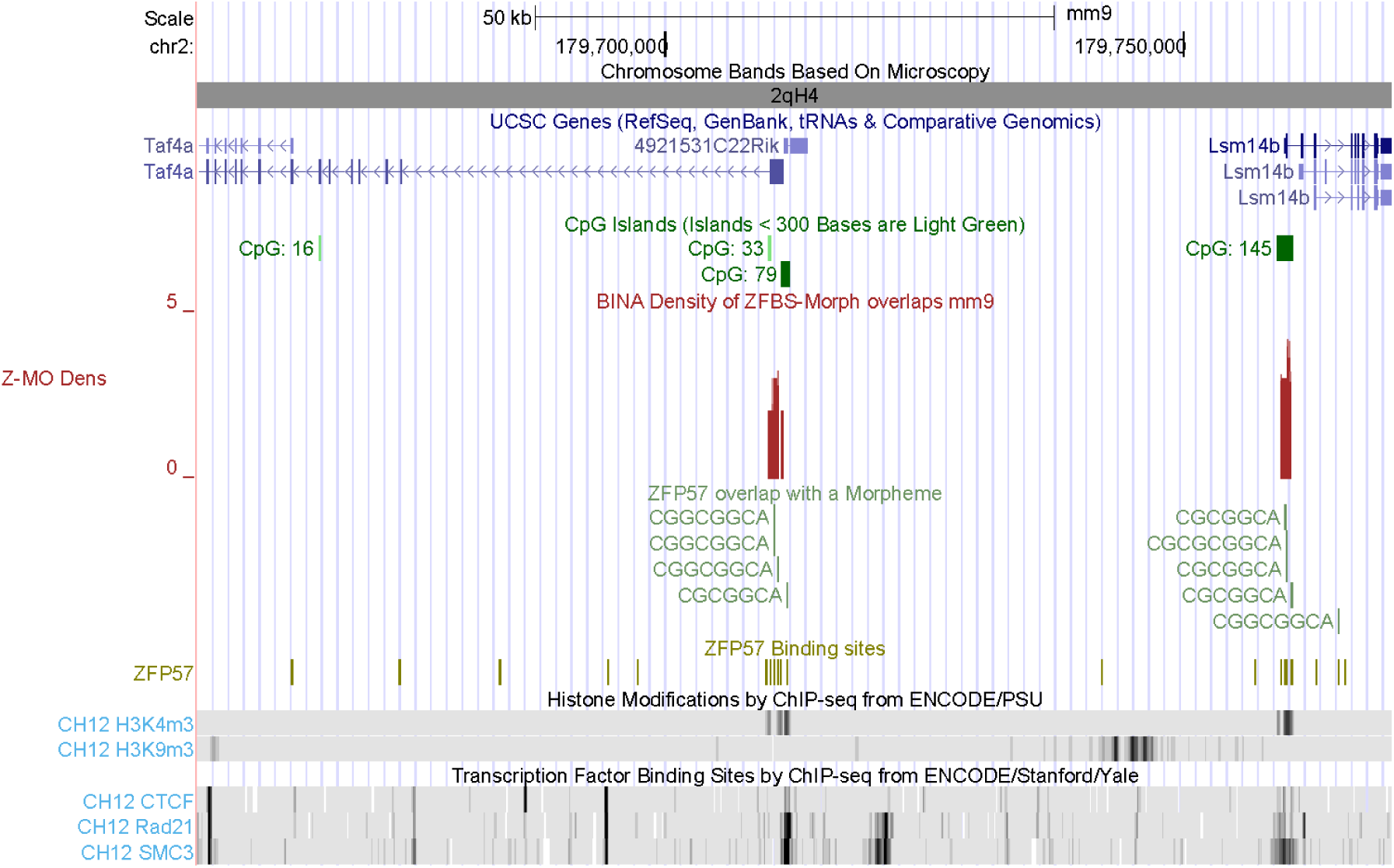
A close-up view of the *Taf4a* and Lsm14b loci. A candidate ICR maps to the longest *Taf4a* transcript, another one maps to the longest *Lsm14b* transcript. A short DNA segment includes the TSSs of both *Taf4a* and *4921531C22Rik* (a testis-specific gene).

**Fig. S2.**
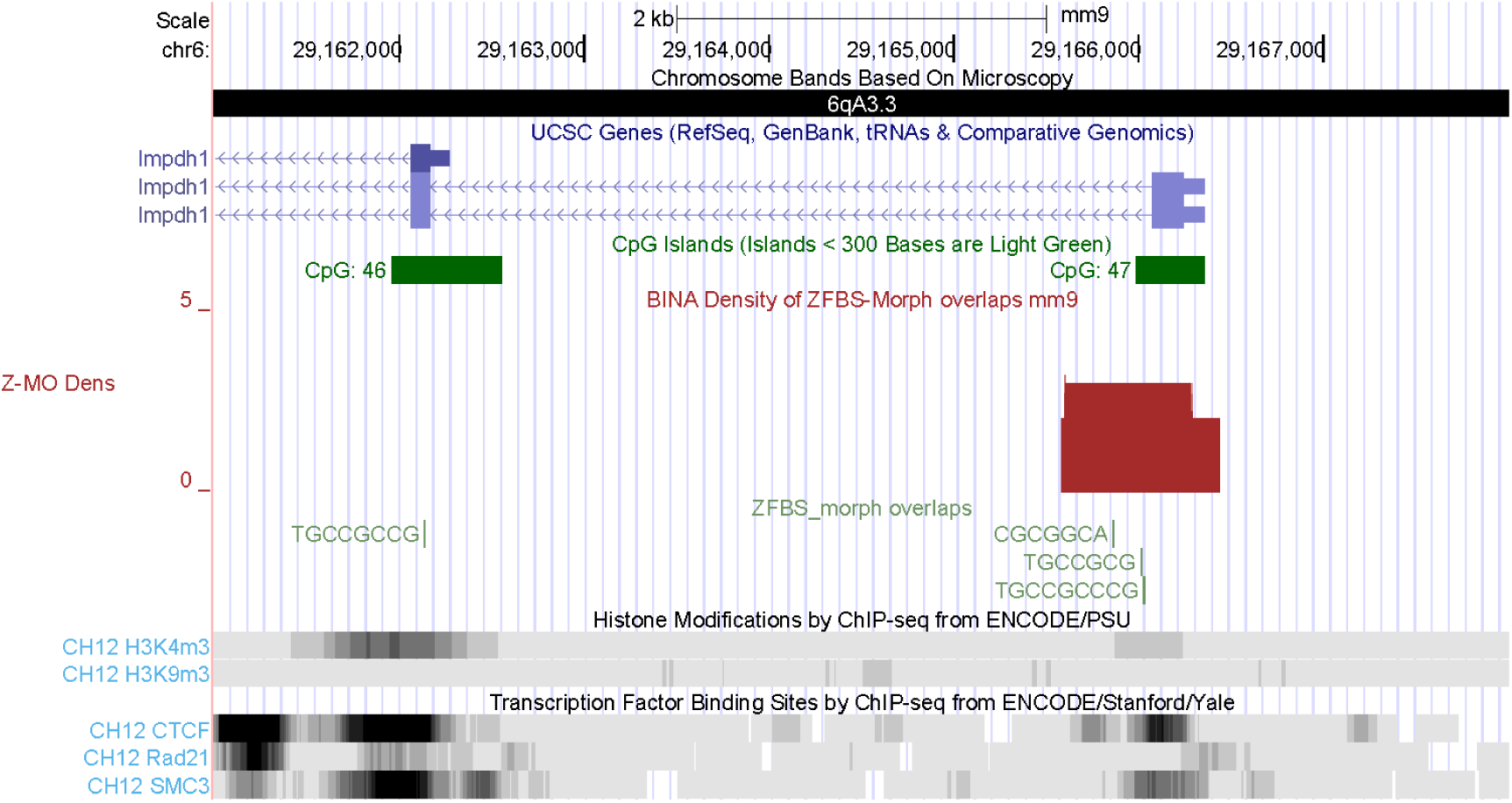
The position of a candidate ICR with respect to the longest *Impdh1* transcripts. The corresponding density peak maps to a CTCF-associated region.

**Fig. S3.**
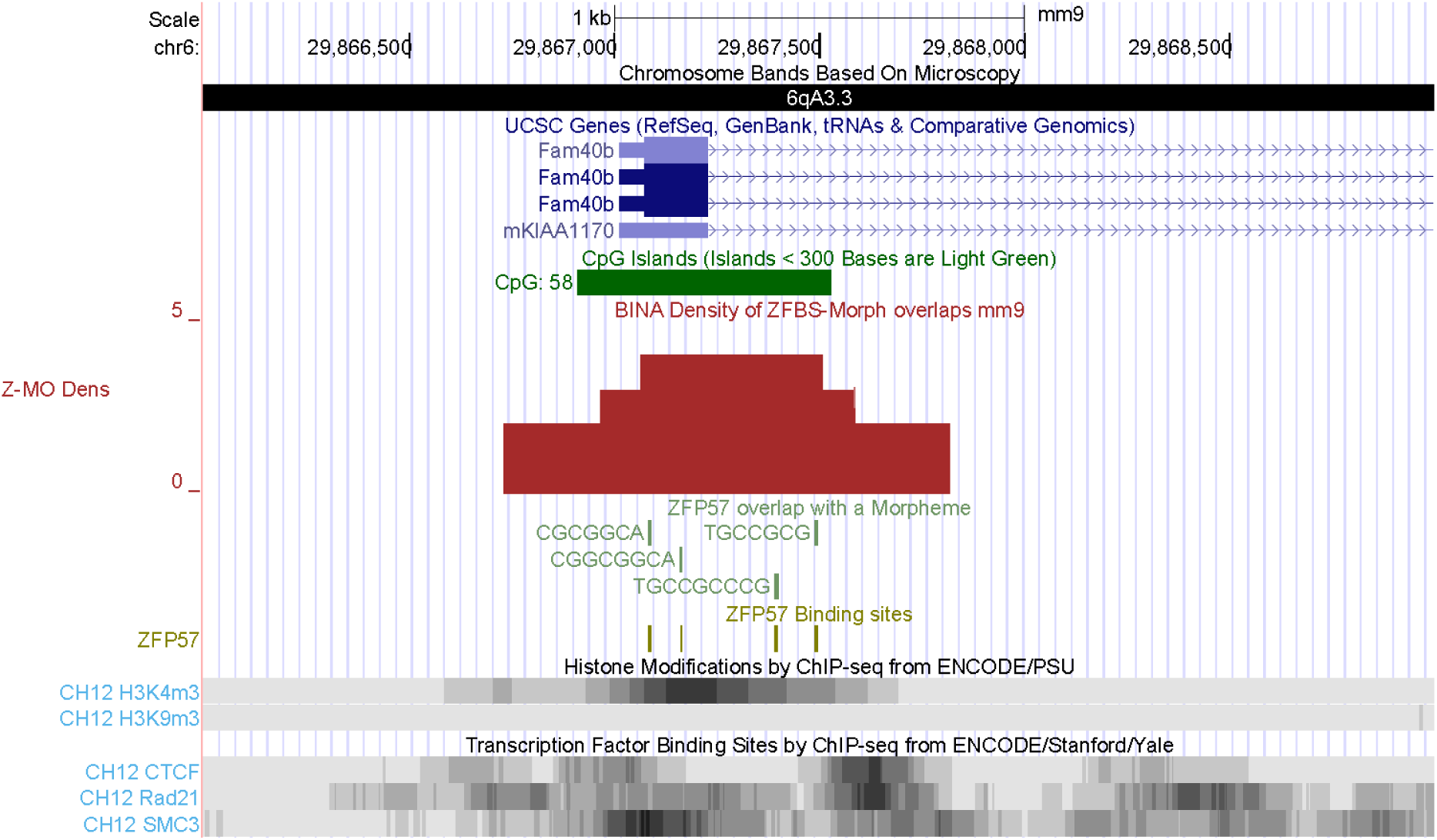
The position of a candidate ICR with respect to the TSS of *Fam40b/ Strip2*. The product of this gene plays an indispensable role in the onset of ESCs differentiation [26].

**Fig. S4.**
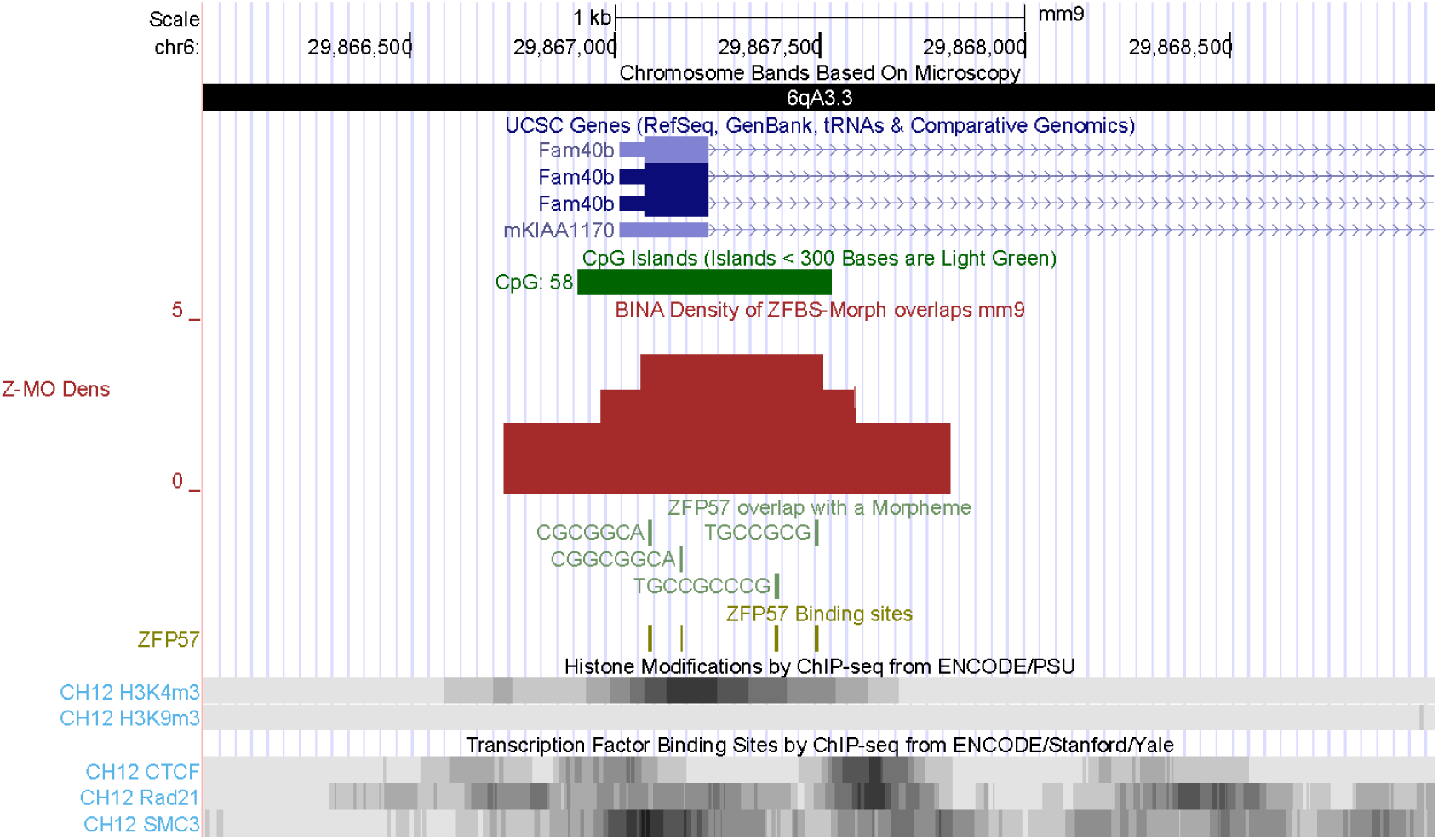
The position of a candidate ICR far upstream of *Slc35b4*. The product of this gene regulates obesity and glucose homeostasis [27]. The displayed segment includes several dispersed H3K9me3 marks.

**Fig. S5.**
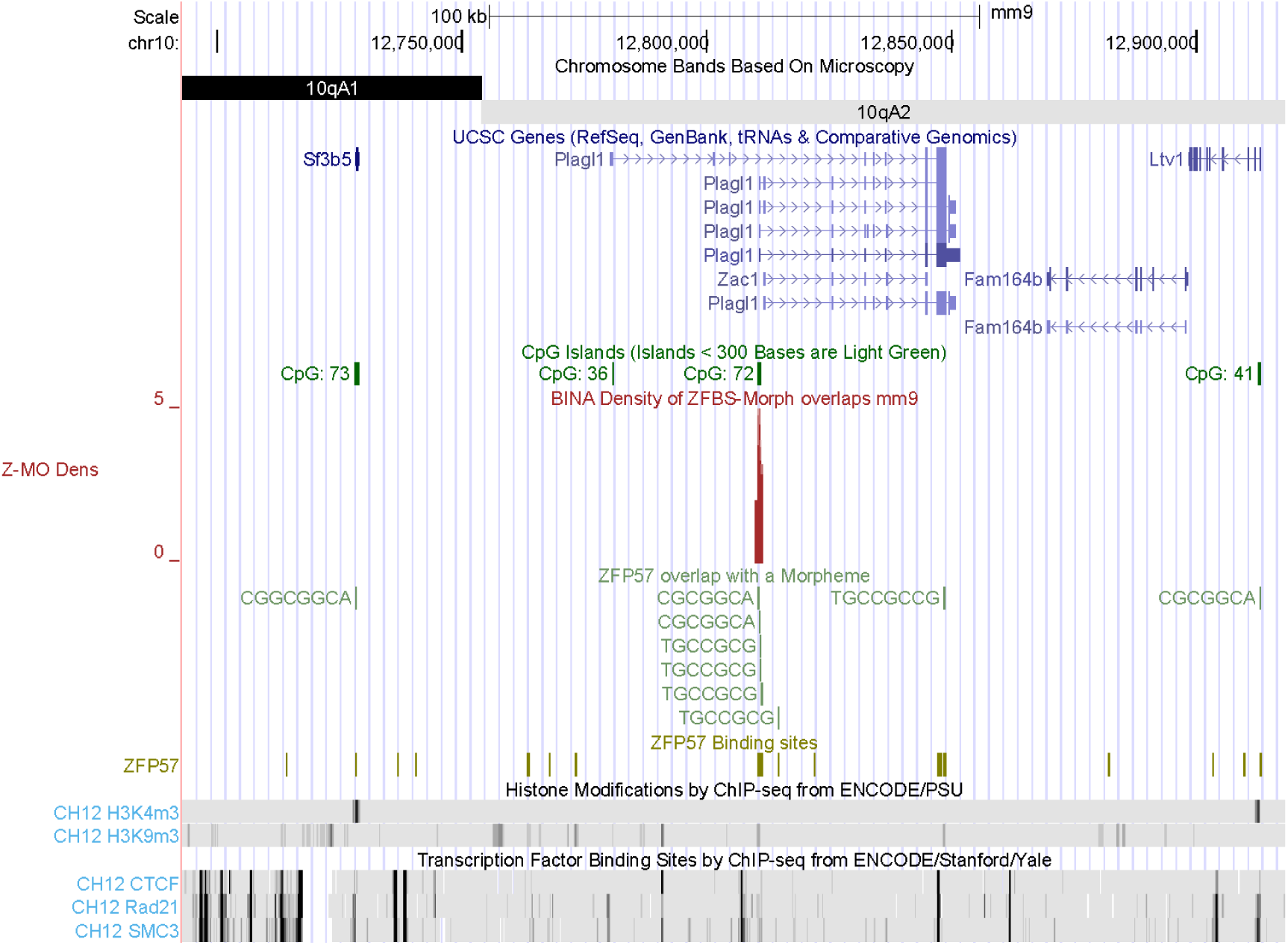
The position of the density peak in the *Plagl1* locus. *Zac1* corresponds to one of the *Plagl1* transcripts. The locus includes a conserved intragenic CpG island that is methylated in oocytes [28]. A very robust density peak correctly located the ICR in the intragenic CpG island (CpG72). This peak covers the previously reported cluster of 5 ZFBS-Morph overlaps in the locus [16].

**Fig. S6.**
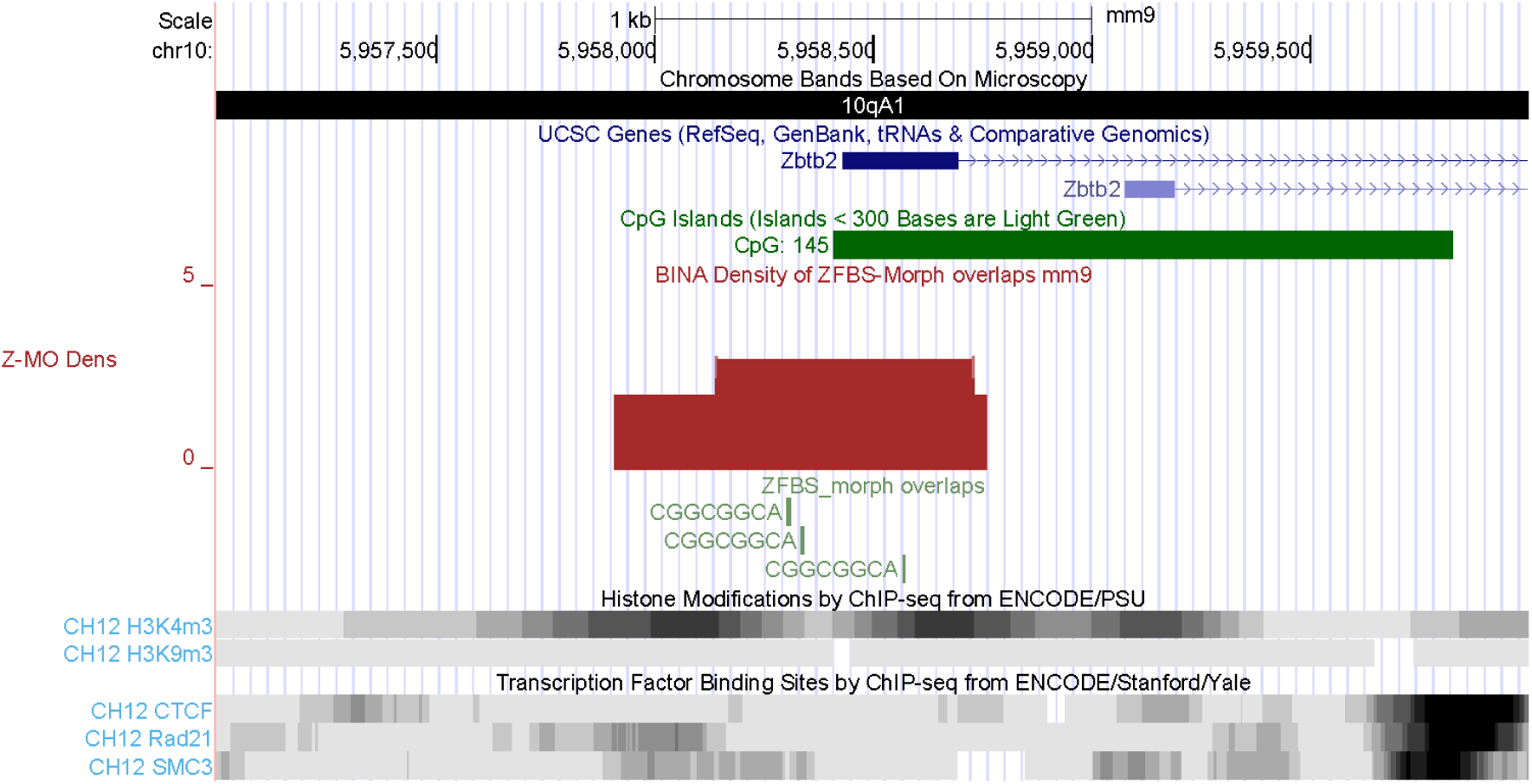
The position of a candidate ICR with respect to the longest *Zbtb2* transcript. *Zbtb2* regulates the differentiation of mouse ES cells [29].

**Fig. S7.**
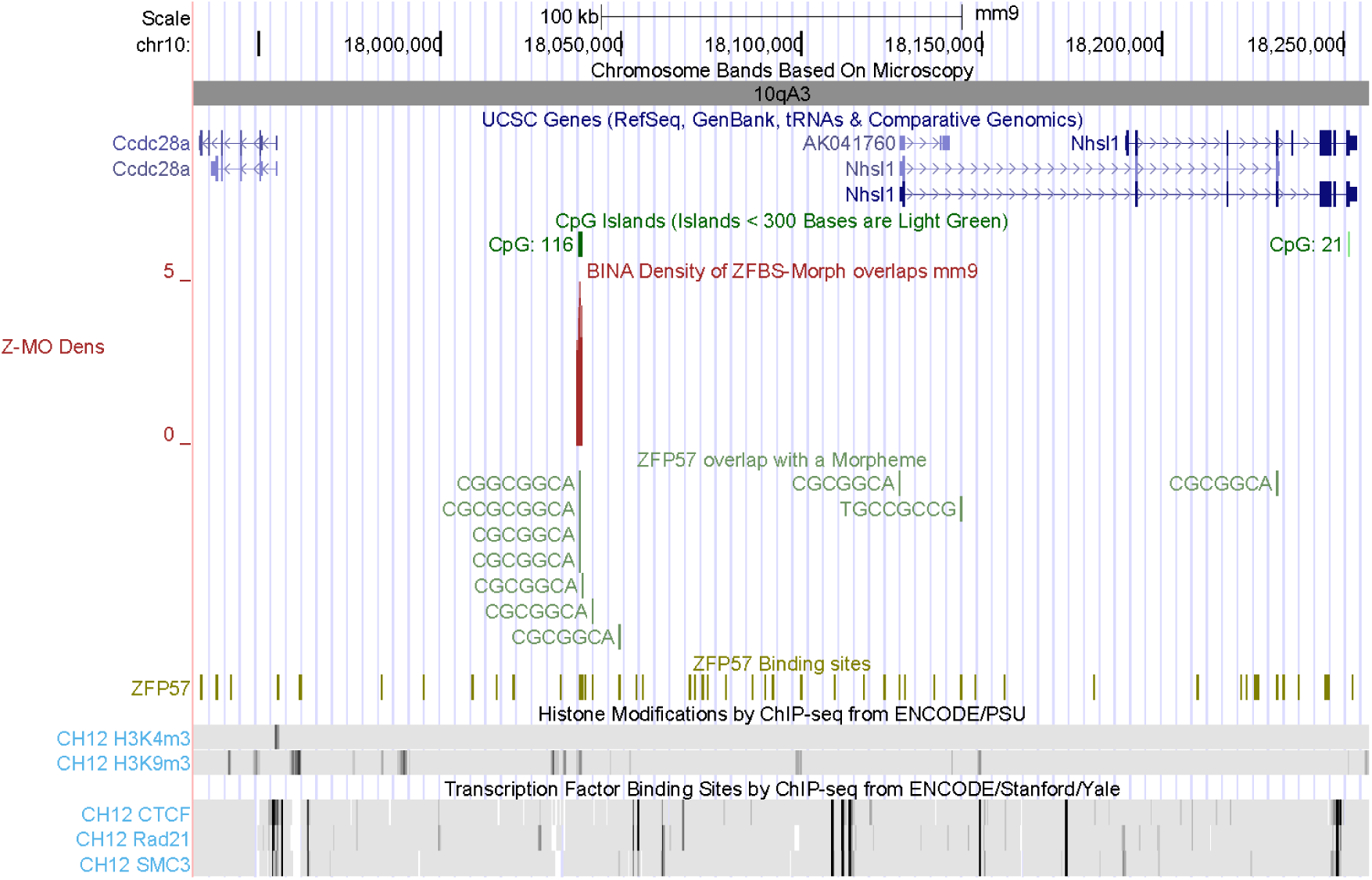
The position of an intergenic candidate ICR with respect to the TSSs of *Ccdc28a* and *Nhsl1. Ccdc28a* is expressed in testes (https://www.ncbi.nlm.nih.gov/gene/215814#gene-expression). The function of *Nhsl1* is not fully understood. The displayed segment includes several dispersed H3K9me3 marks.

**Fig. S8.**
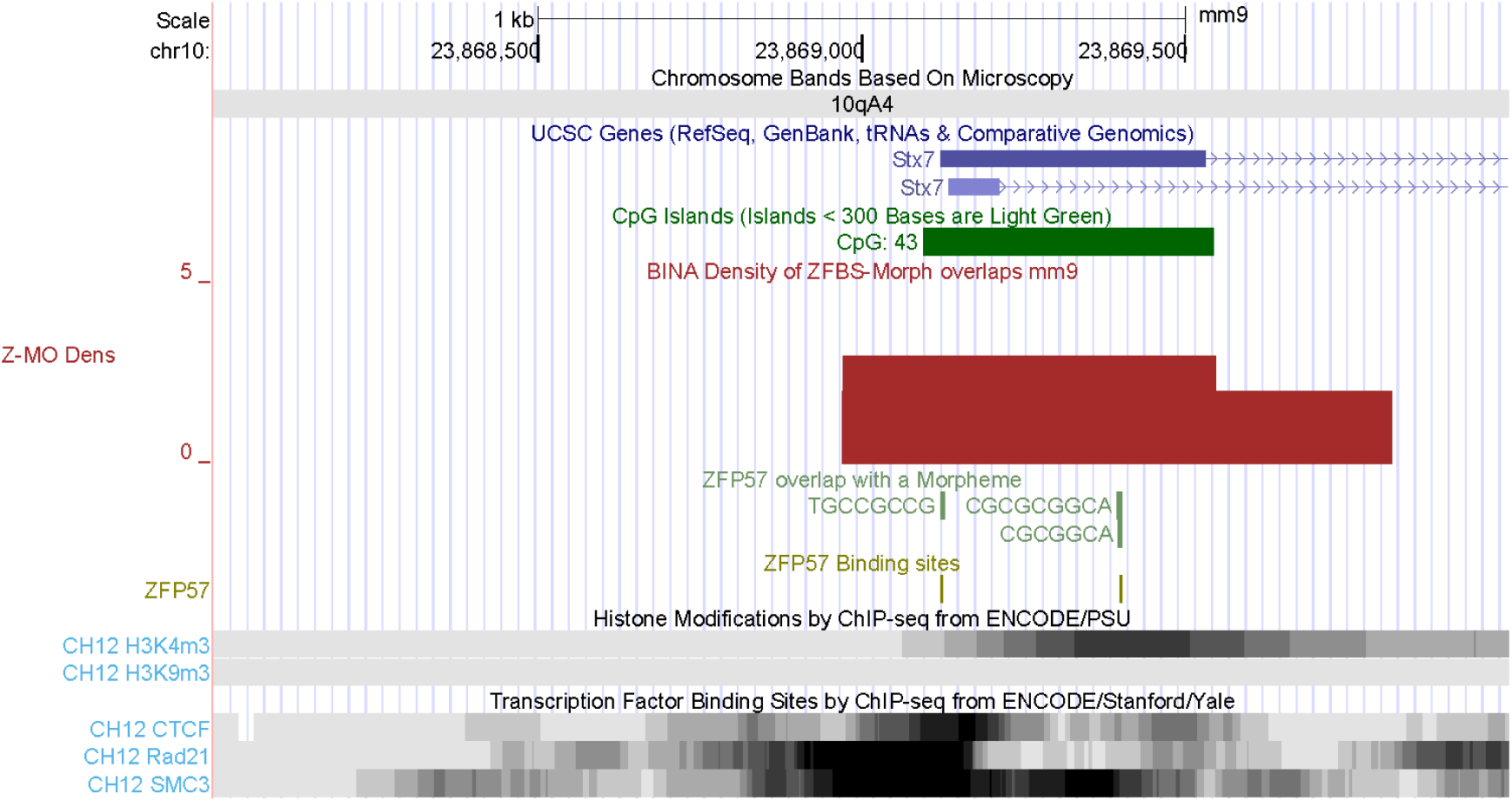
The position of a candidate ICR within the *Stx7* locus. The corresponding density peak overlaps a chromatin boundary consisting of CTCF, RAD21, and SMC3. Syntaxin-7 is among a group of proteins that function in vesicle transport and fusion events [30].

**Fig. S9.**
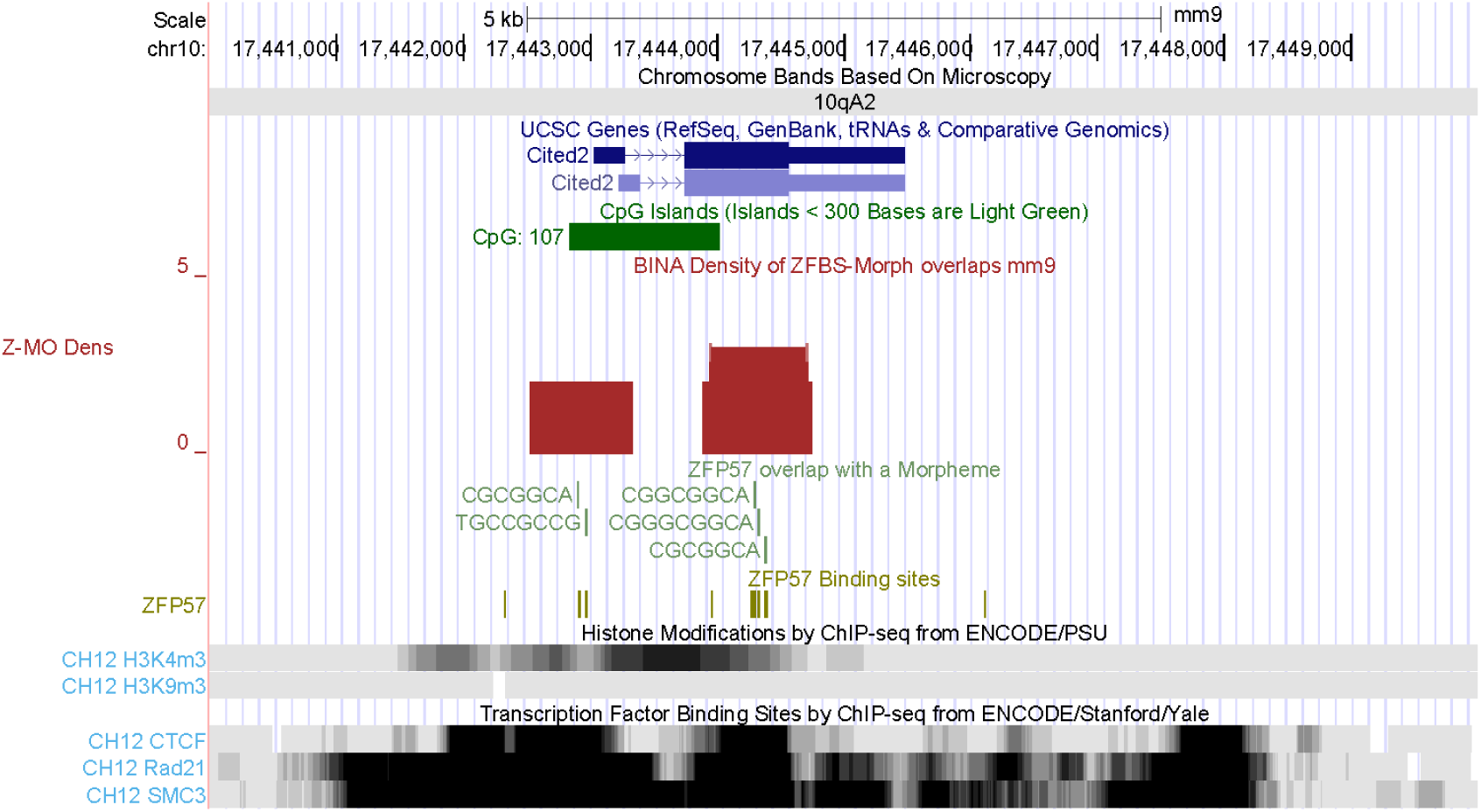
The position of a candidate ICR in the *Cited2* locus. This ICR contains 2 density peaks that map to chromatin boundaries. Absence of *Cited2* in mouse embryos caused congenital heart disease by perturbing left-right patterning of the body axis [33]. The locus includes several closely-spaced chromatin boundaries.

**Fig. S10.**
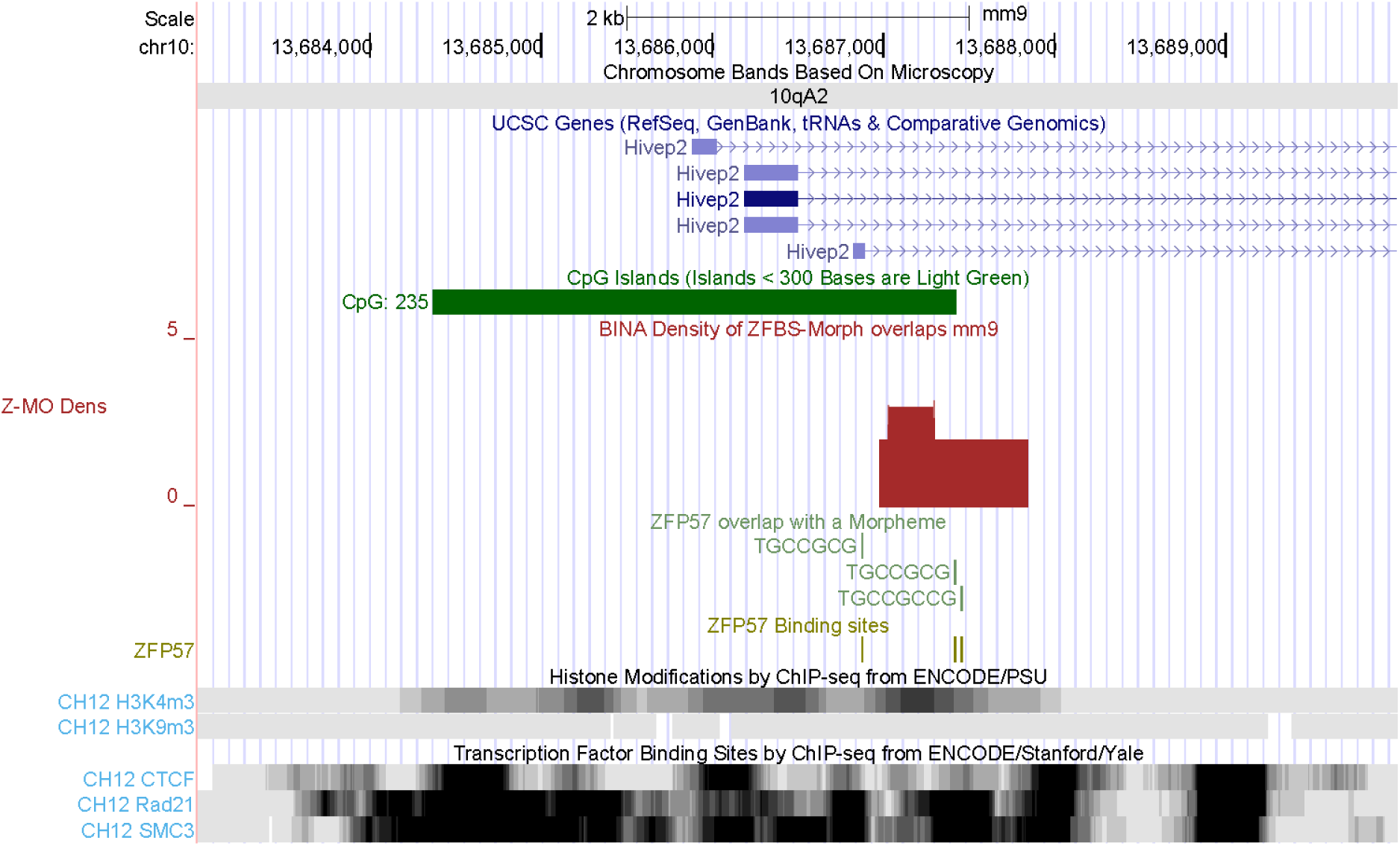
An intragenic candidate ICR within the *Hivep2* locus. The corresponding density peak is in a region that encompasses several chromatin boundaries. *Hivep2* deficiency conferred molecular, neuronal, and behavioral phenotypes related to schizophrenia [32].

**Fig. S11.**
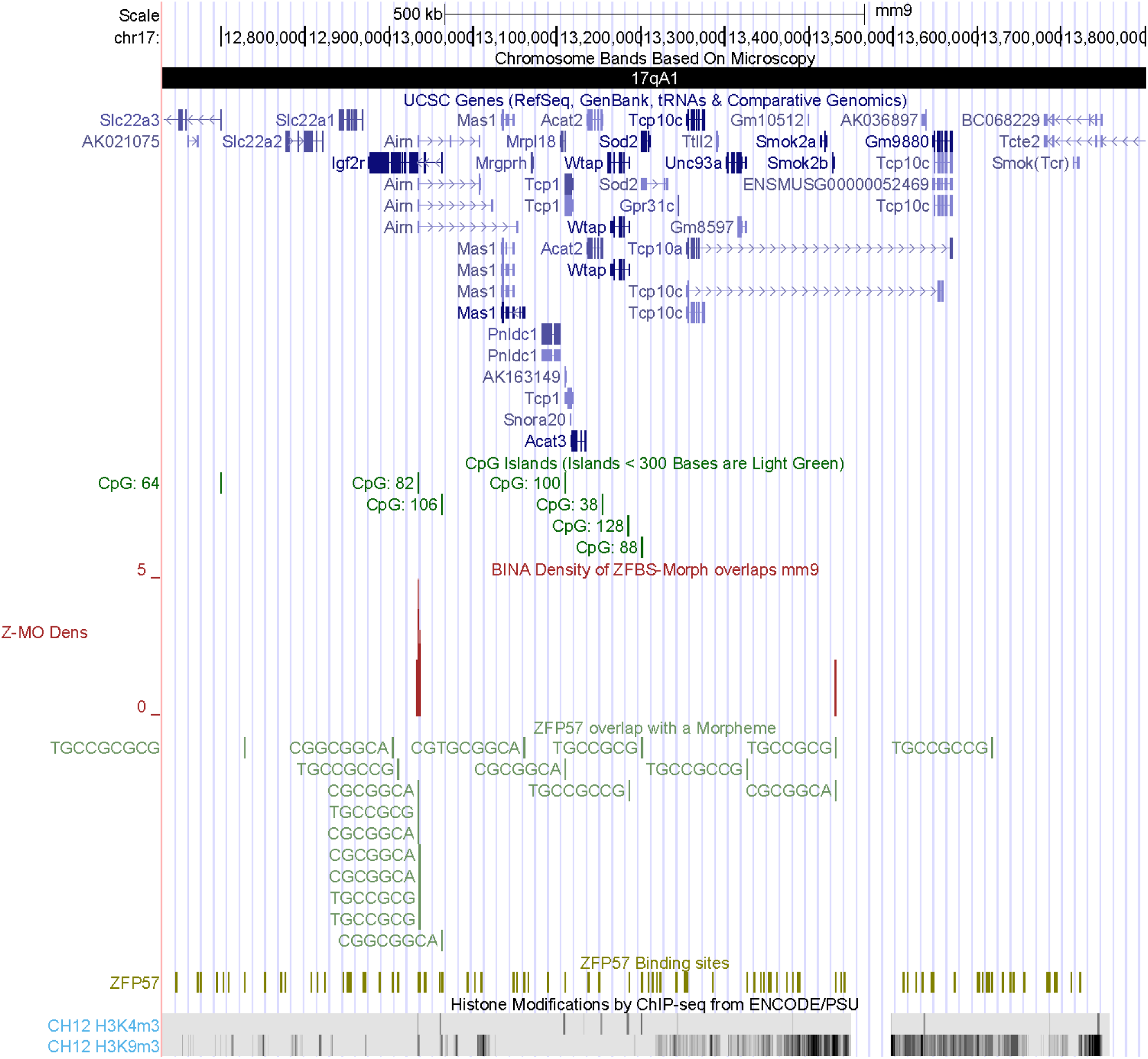
The position of the density peak in the *Igf2r* – *Airn* imprinted domain. This peak covers several ZFBS-morph overlaps located in the intronic CpG island that regulates expression of genes in the imprinted domain [16]. The displayed DNA section consists of 1.17 Mb. This section also includes *Tcp1* –one of the genes carried by all t haplotypes [93]. Note that there is a density peak in the longest intron of Tcp10. This gene also is in the chromosomal section that encompasses the t haplotypes. Even though the peak covers only 2 ZFBS-morph overlaps, it is within a relatively long chromatin segment that includes extensive repressive H3K9me3 marks.

**Fig. S12.**
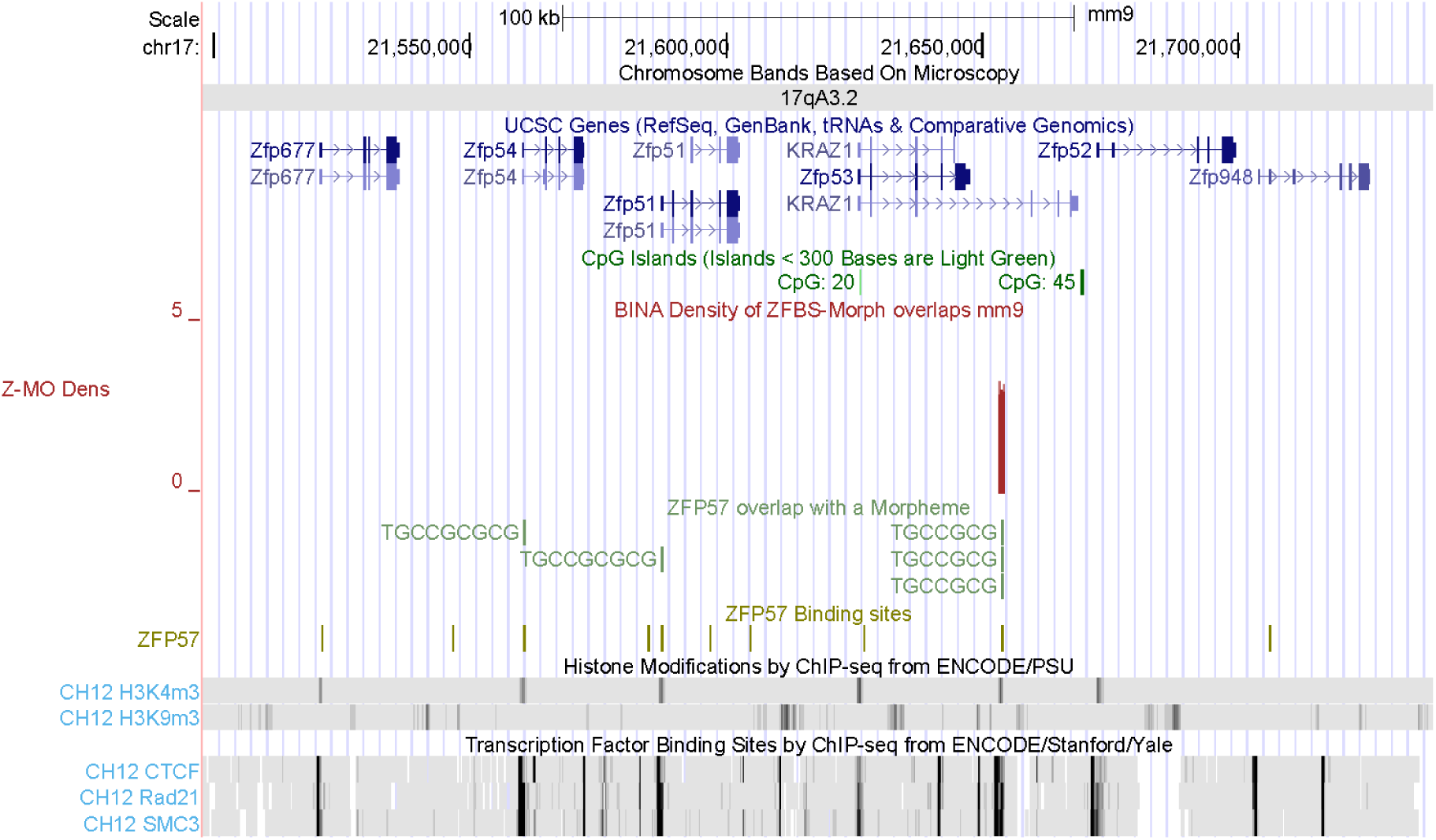
The position of a candidate ICR for a gene (*KRAZ1*) in a chromosomal section with several ZFP genes. *KRAZ1* encodes a factor with Krüppel-type zinc finger structural motif [42-44]. As ZNF57, KRAZ1 interacts with KAP1/TRIM28 [43]. The displayed segment includes several dispersed repressive H3K9me3 marks.

**Fig.S13.**
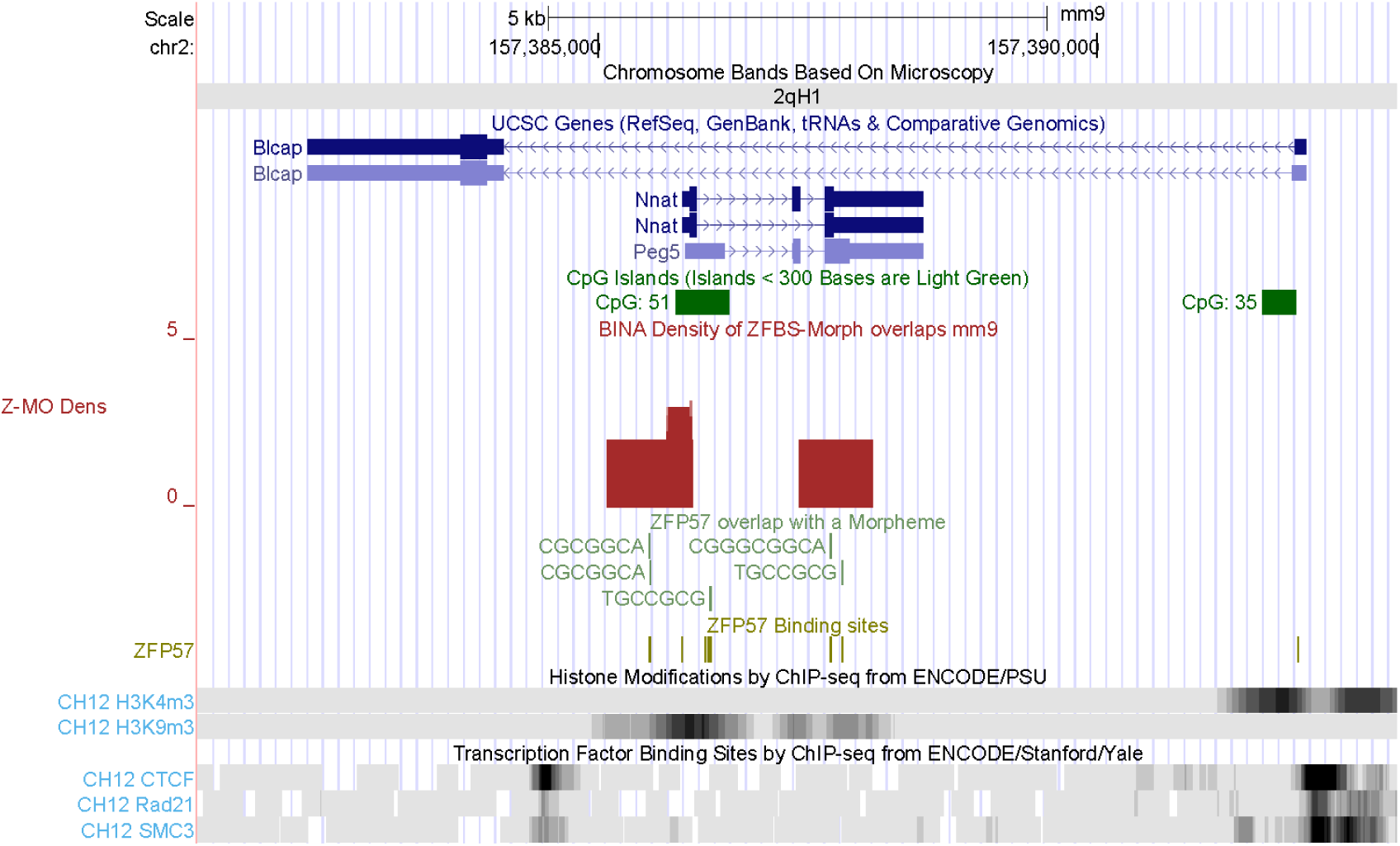
Two intragenic density peaks locating the ICR of *Nnat* in *Blcap* locus.

**Fig. S14.**
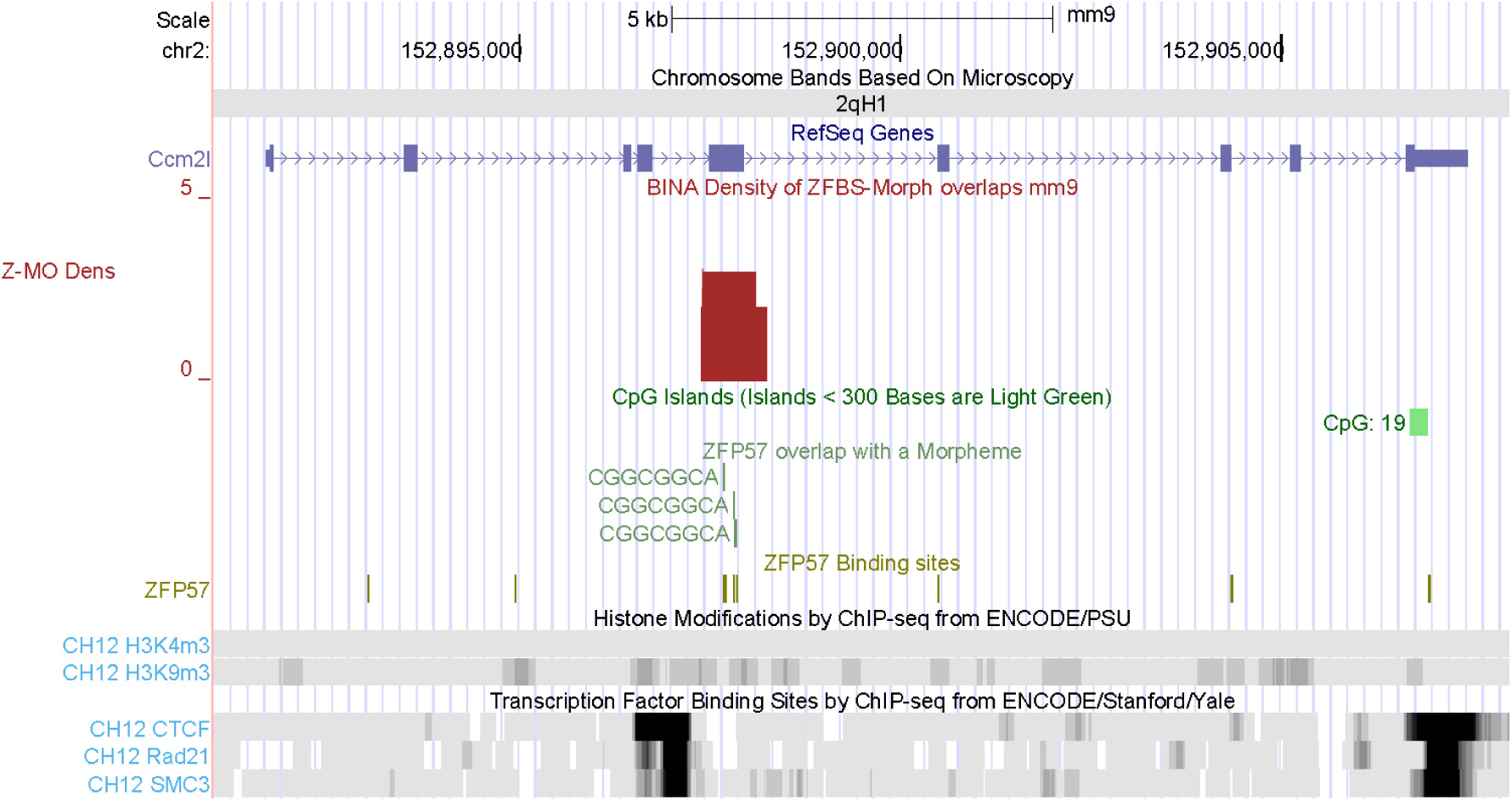
A candidate ICR mapping to *Ccm2l* locus. The product of this gene is selectively produced in endothelial cells during angiogenesis [57]. *Ccm2 l*^-/-^ animals exhibited embryonic lethality at E11 associated with myocardial thinning [56].

**Fig. S15.**
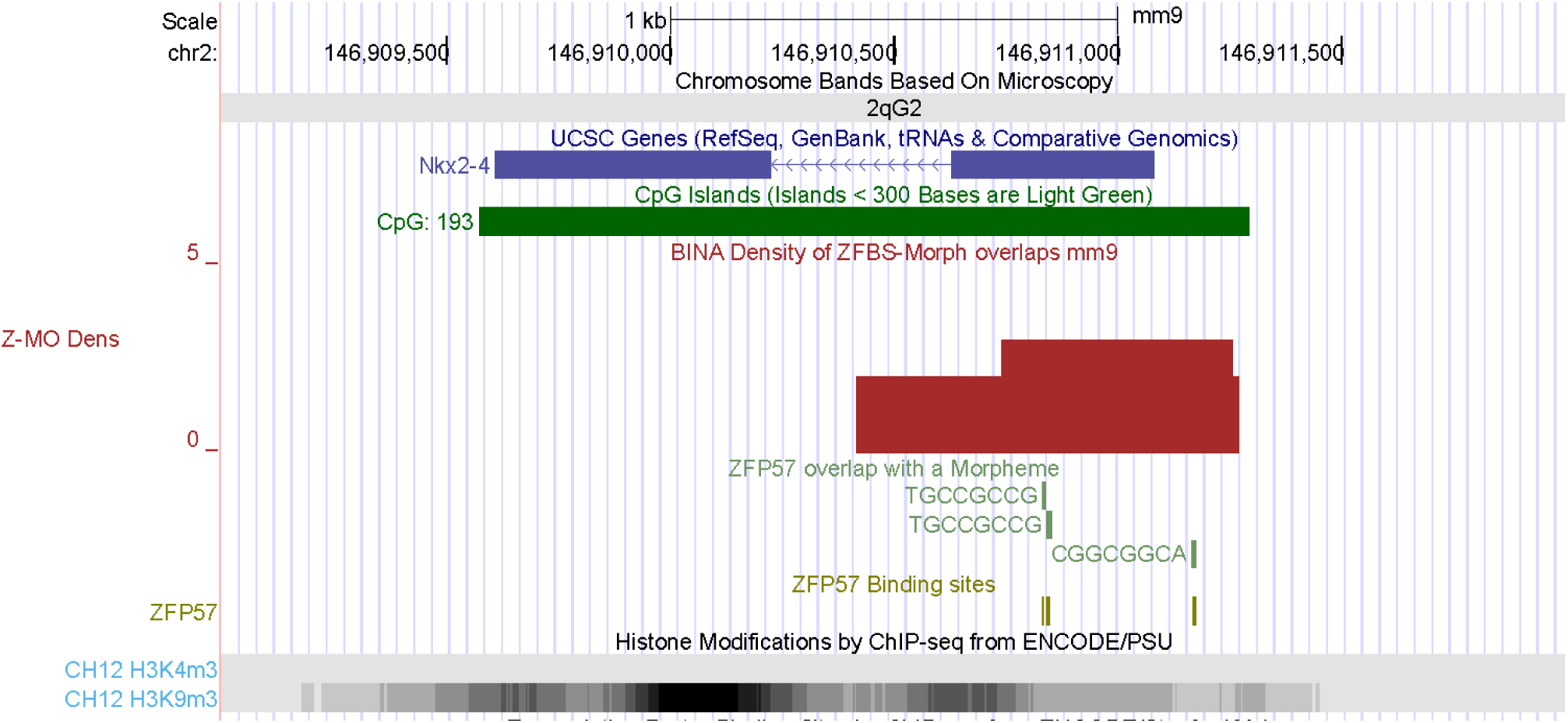
The position of a candidate ICR mapping to *Nkx2-4.* The product of this gene is a transcription factor whose structure includes a homeodomain for binding DNA [58]. During mouse embryogenesis, *Nkx-2.2* transcripts were dispersed in localized domains of the brain and might be involved in specifying diencephalic neuromeric boundaries [58]. The displayed segment includes dispersed H3K9m3 marks.

**Fig. S16.**
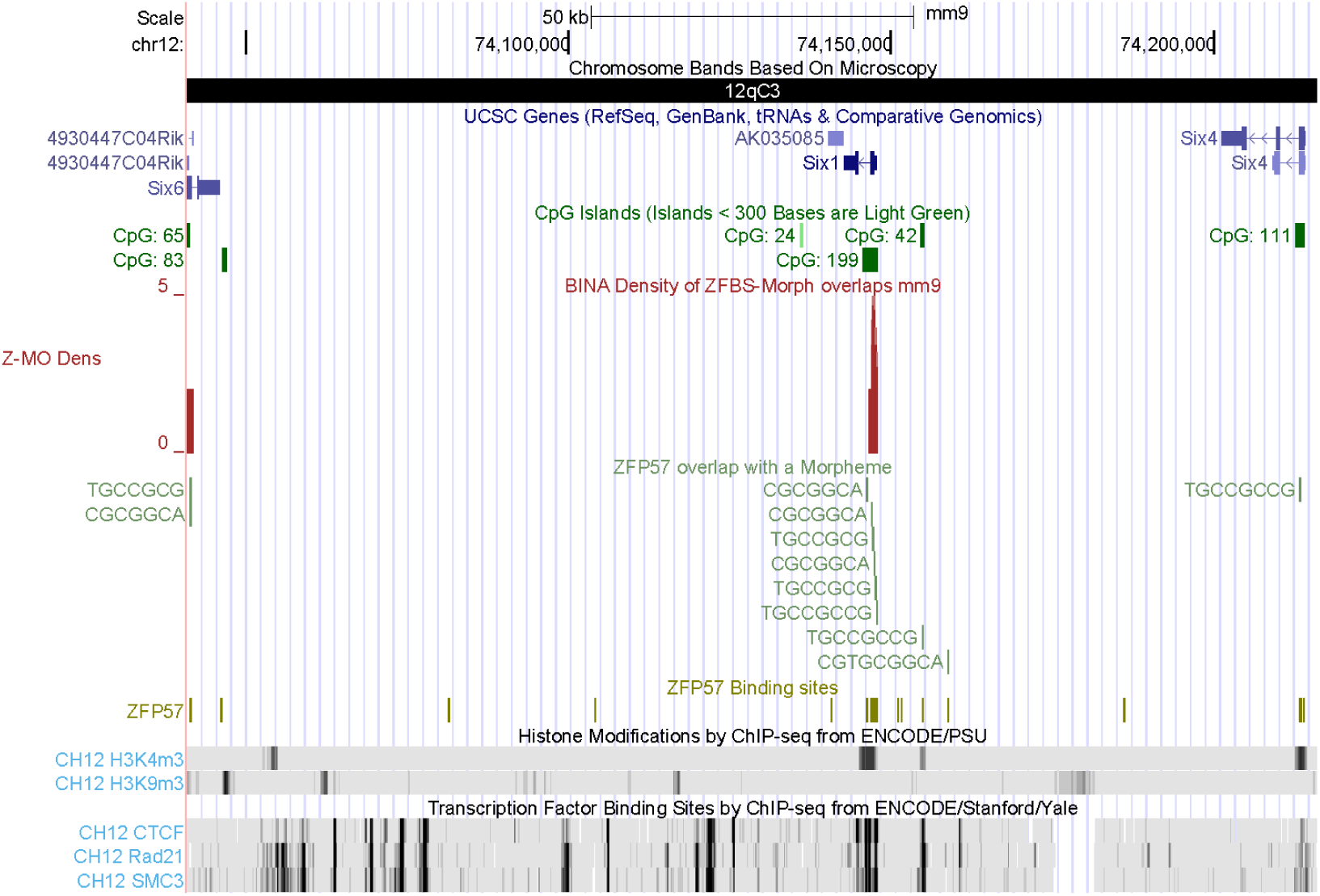
A candidate ICR in a DNA segment with several *Six* genes. The corresponding density peak maps to *Six1*. This gene specifies a transcription factor (SIX1) that is a component of a protein-network that controls organ development [62]. *Six1*-deficient mice embryos were devoid of inner ear structures [94].

**Fig. S17.**
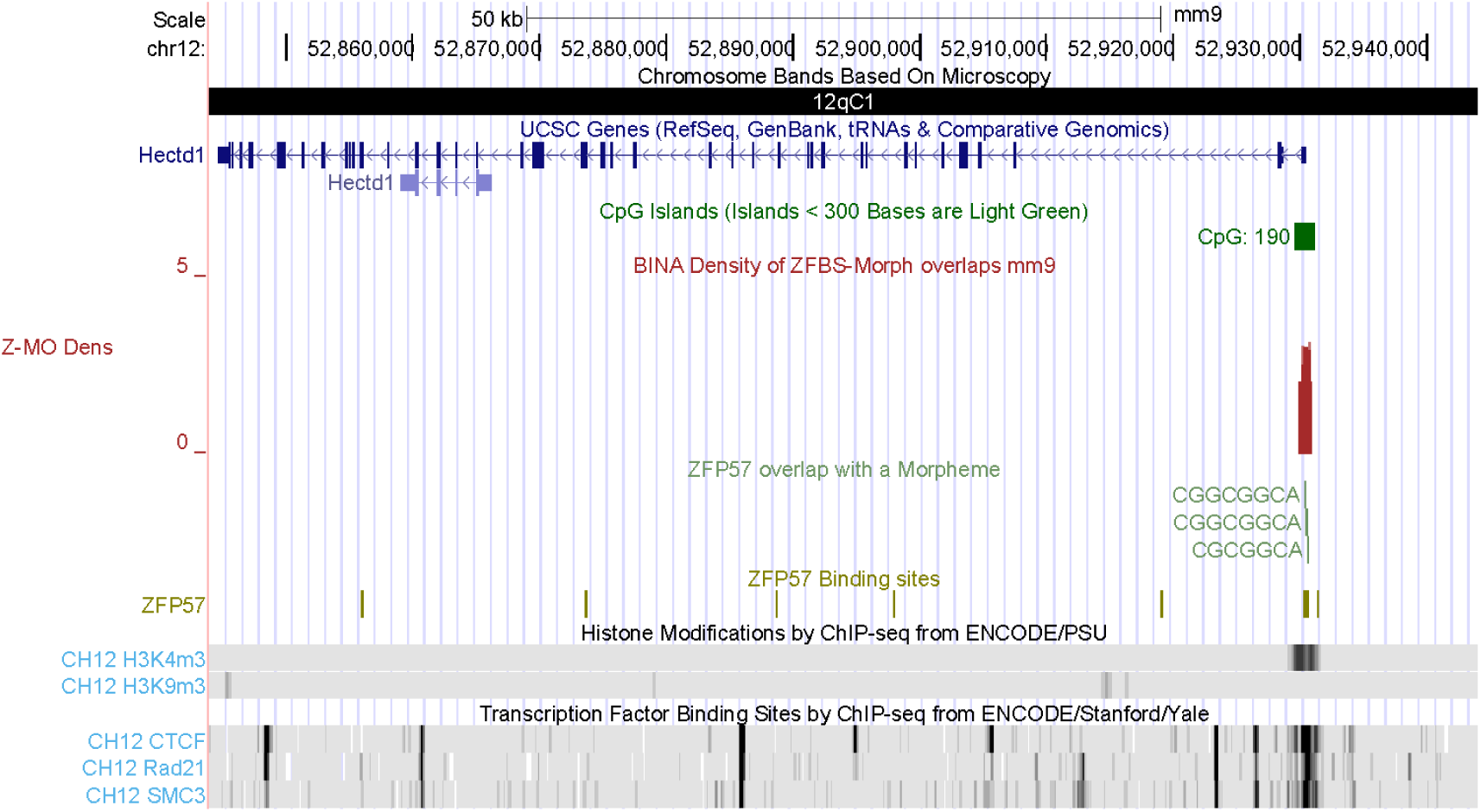
The position of a candidate ICR mapping to *Hectd1* locus. The corresponding density peak is within chromatin boundaries. Disruption of *Hectd1* resulted in mid-gestation lethality and intrauterine growth restriction [63].

**Fig. S18.**
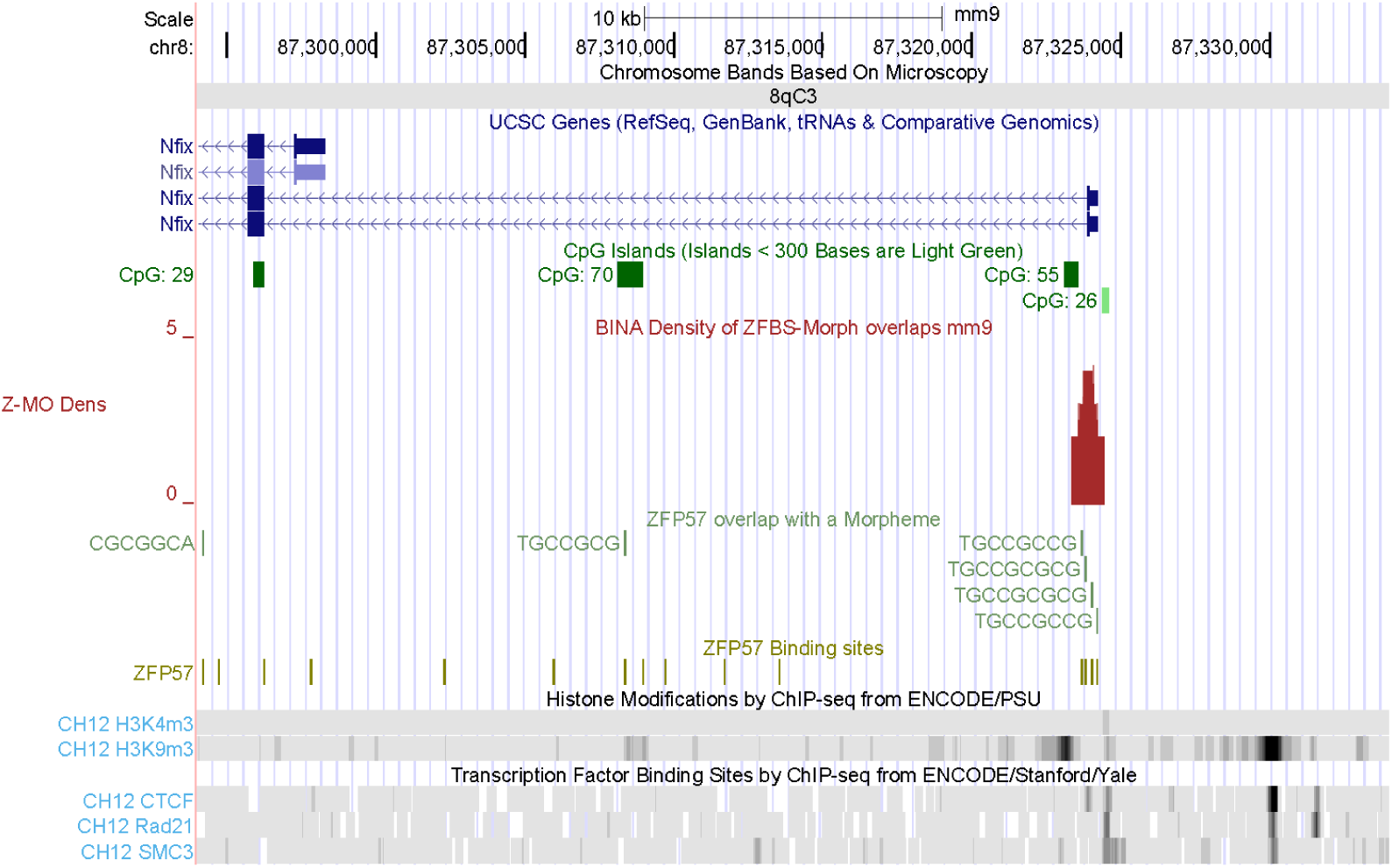
The position of a candidate ICR mapping to the longest *Nfix* transcripts. NFIX belongs to a family of transcription factors that play important roles in the development of the nervous system [66]. The locus includes dispersed H3K9me3 marks.

**Fig. S19.**
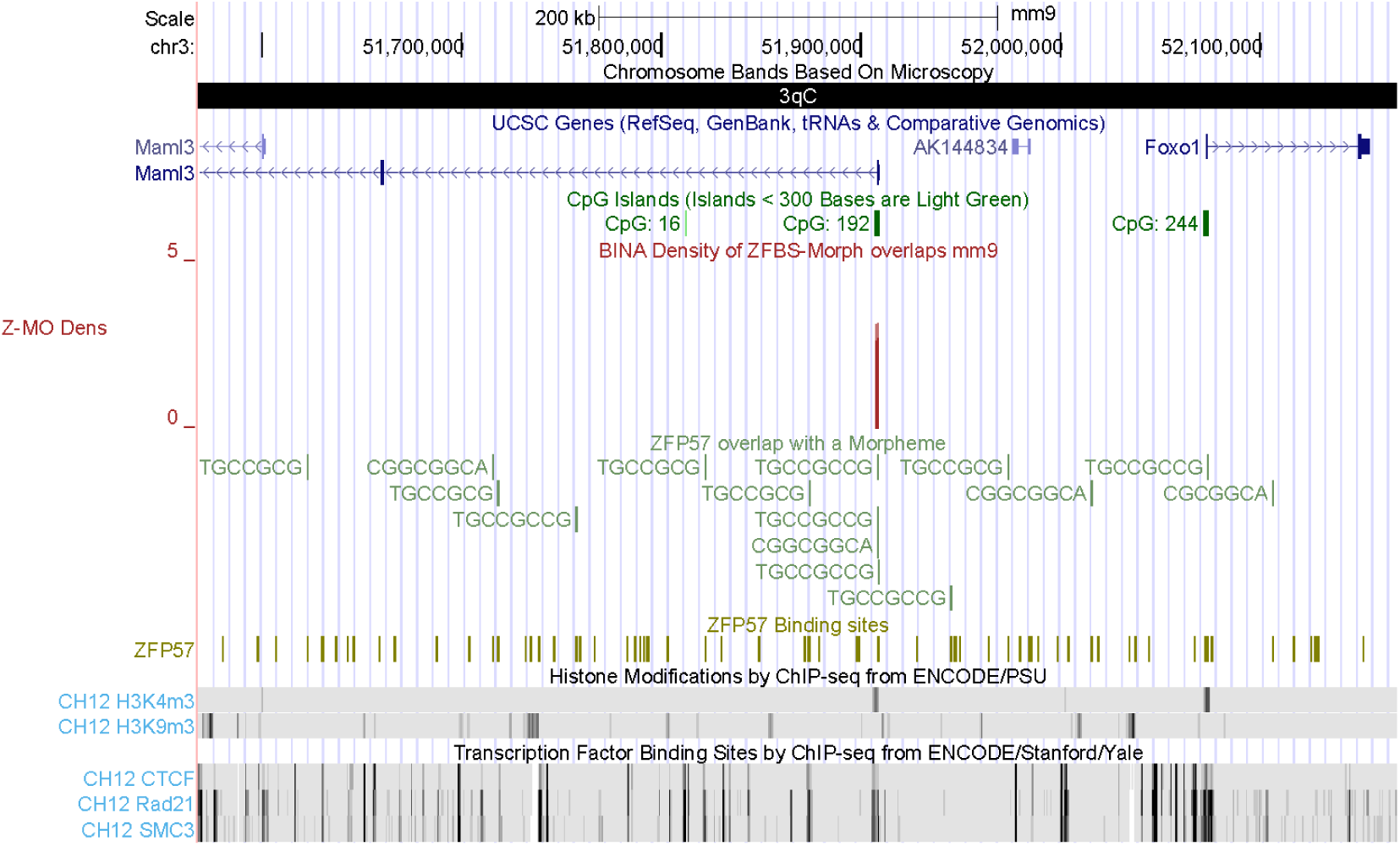
The position of a candidate ICR mapping to the longest of *Maml3* transcript. This candidate ICR is far upstream of *Foxo1* TSS. *Maml3* functions in organogenesis [67]. Across mammals, *Foxo1* functions in both male and female reproductive processes [95].

**Fig. S20.**
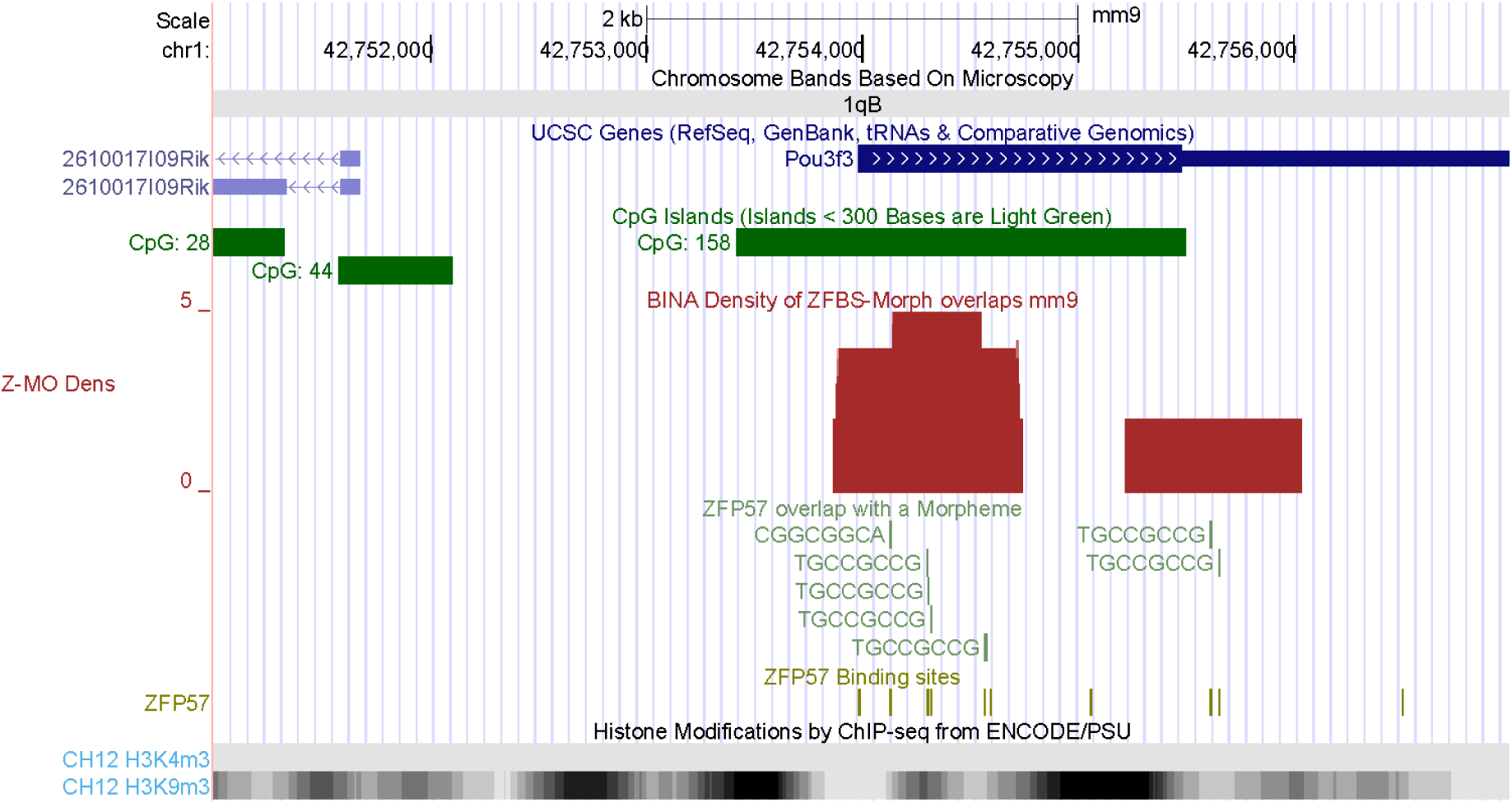
Density peaks defining a candidate ICR for *Pou3f3*. POU3F3 influences neurogenesis [68]. Upstream of *Pou3f3* is the TSS of a noncoding RNA gene (*Pantr1*). This RNA is differentially expressed during neural stem cell differentiation [96]. Dispersed between *Pou3f3* and *Pantr1* are several regions containing repressive H3K9m3 marks.

**Fig.S21.**
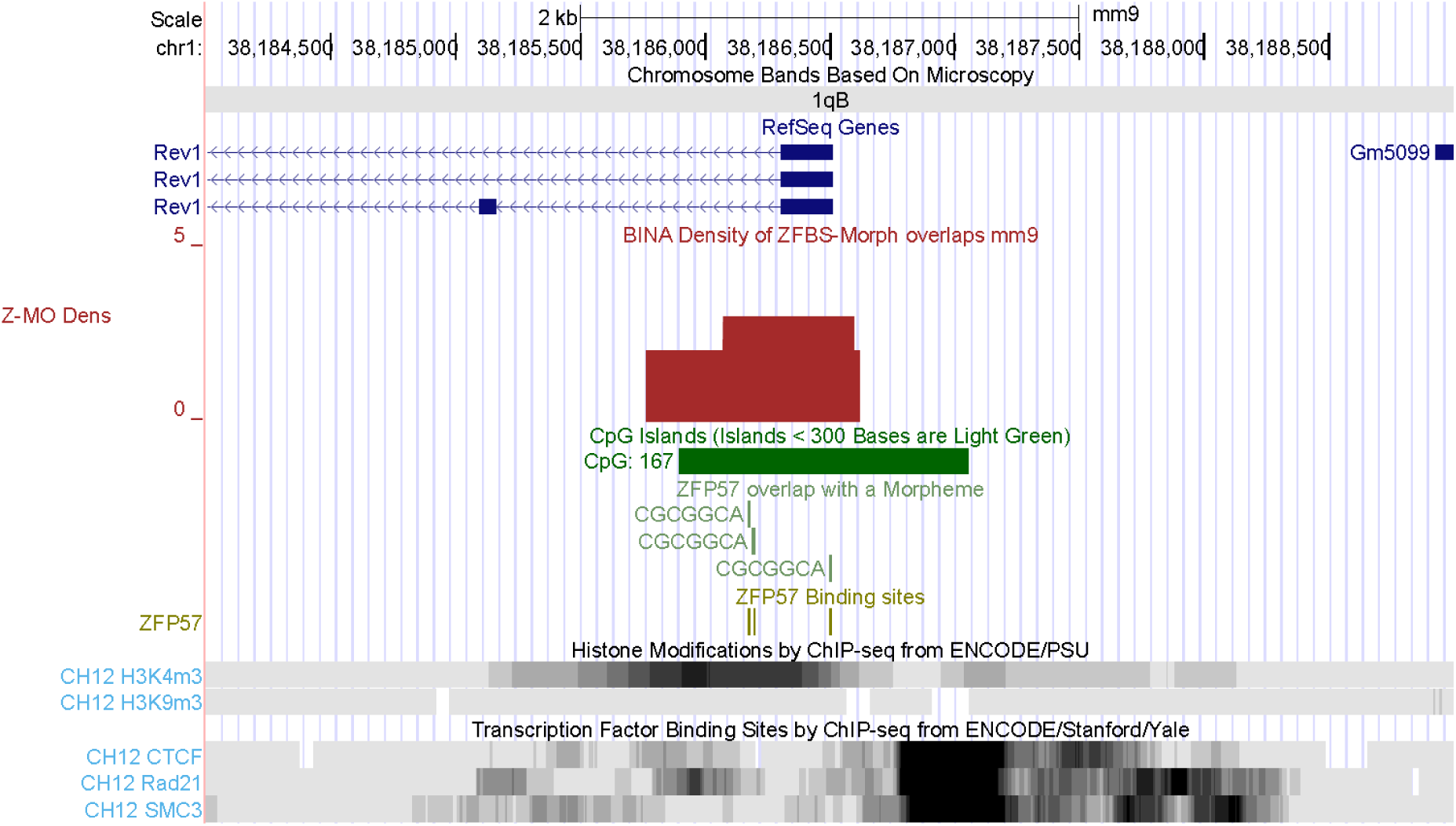
The position of a candidate ICR mapping to *Rev1* locus. This gene encodes a DNA polymerase that during somatic hypermutation incorporates deoxycytidine residues most likely opposite abasic nucleotides [97].

**Fig. S22.**
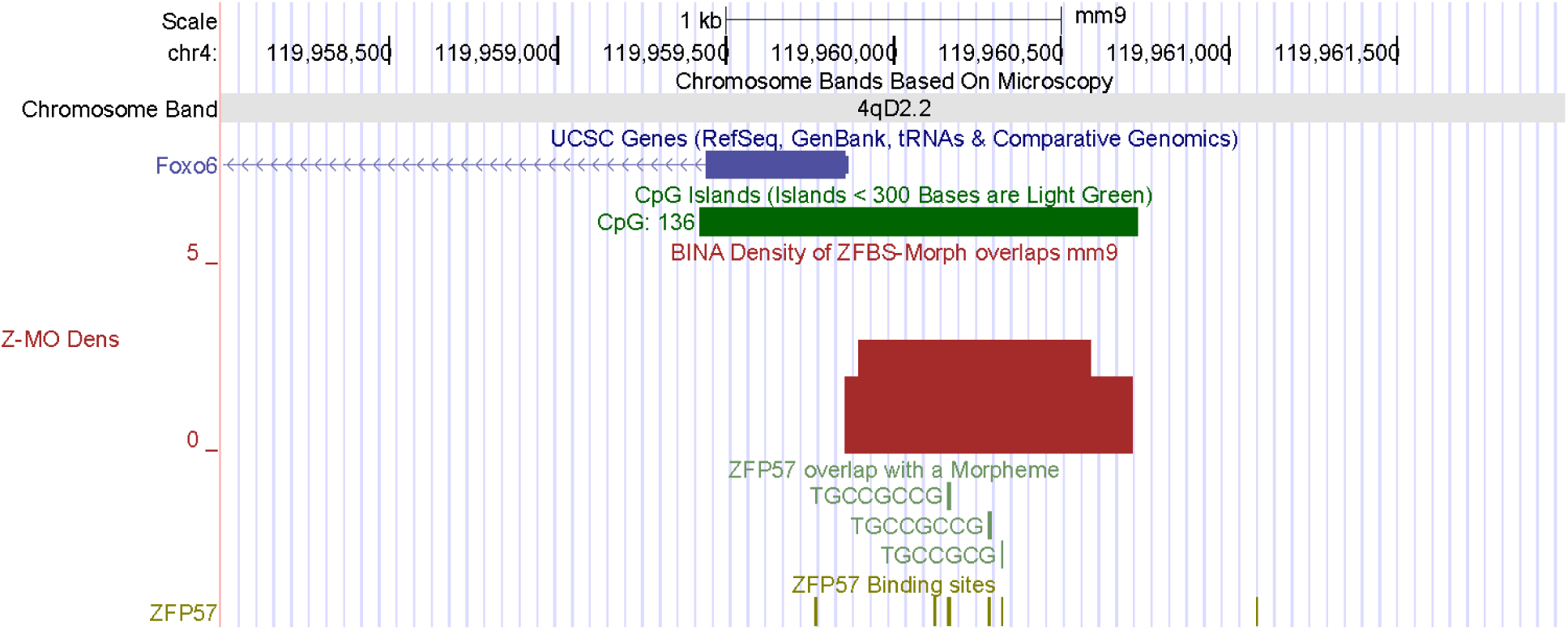
The position of a candidate ICR for *Foxo6.* This gene encodes a transcription factor. Its functions include regulation of Hippo signaling and growth of the craniofacial complex [71].

**Fig. S23.**
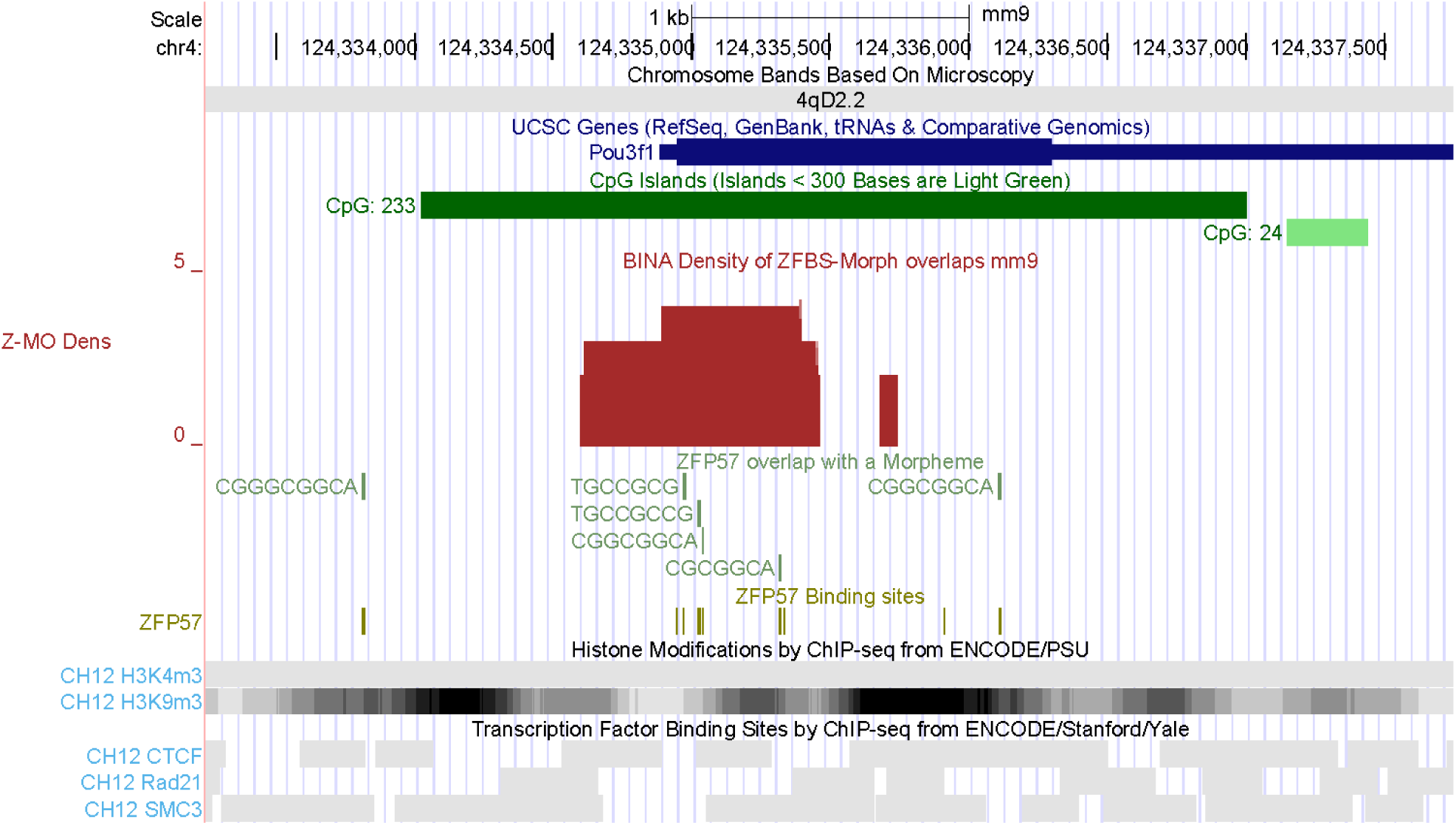
The position of a candidate ICR for *Pou3f1*. A reduction in *Pou3f1* expression induced apoptosis of cultured germ cells [72]. Biochemical studies localized POU3F1 to spermatogonia of prepubertal and adult testes [72].

**Fig. S24.**
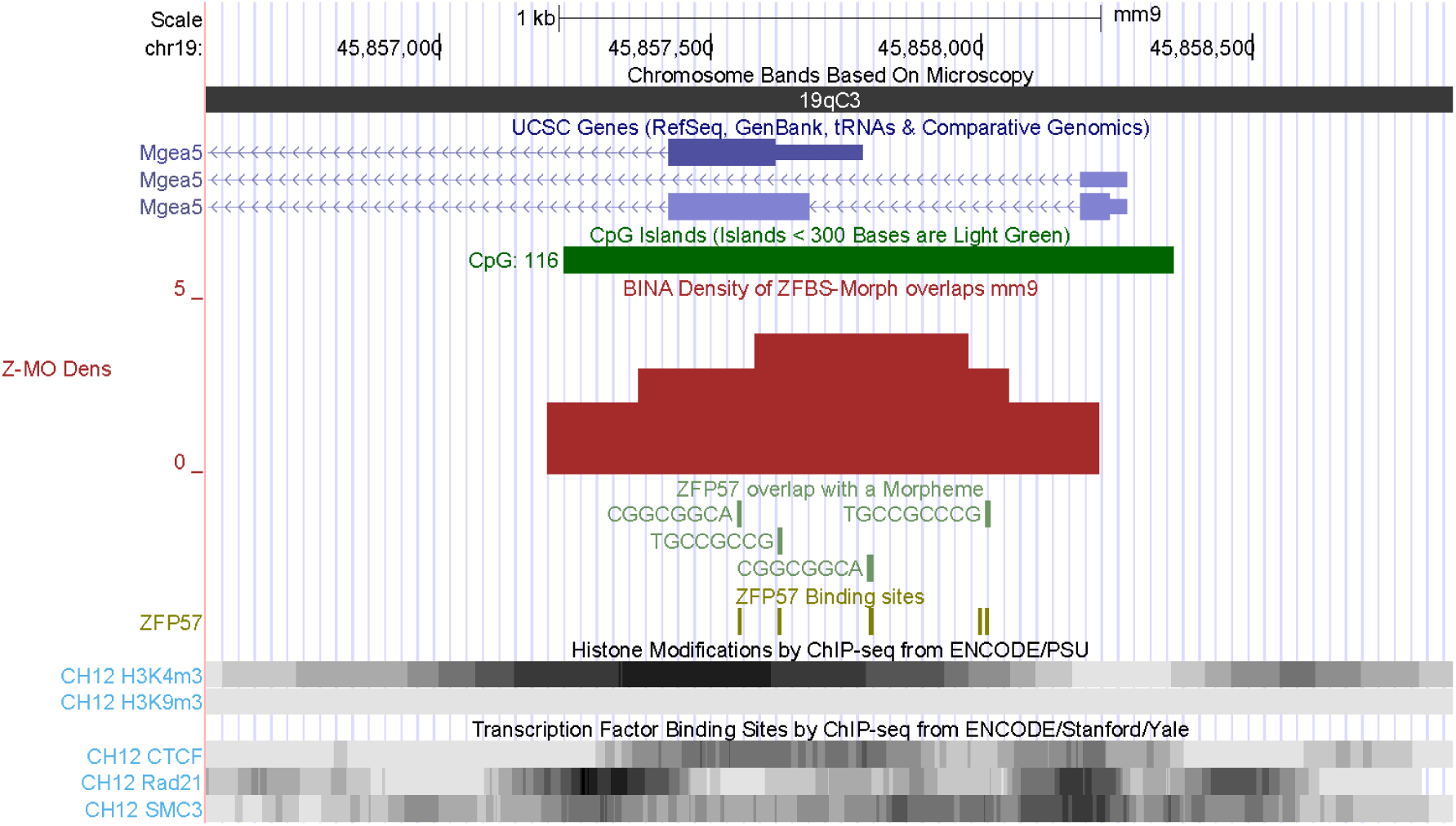
The position of a candidate ICR for the longest *Mgea5* transcript. This gene encodes an enzyme that influences folliculogenesis [73].

**Fig. S25.**
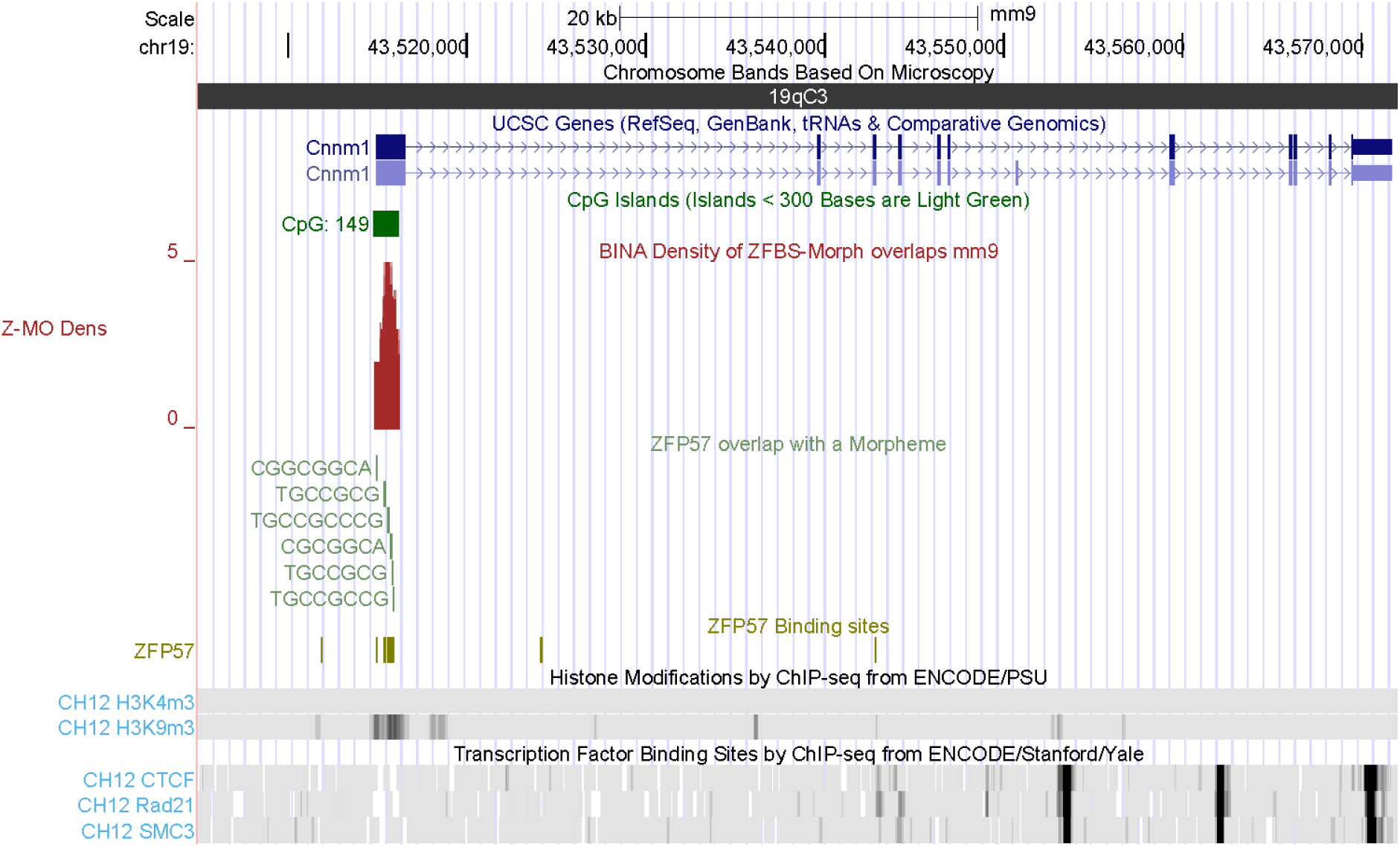
The position of a candidate mapping to *Cnnm1*. Expression of this gene is associated with cell cycle and differentiation in spermatogenic cells in mouse testis [74].

**Fig. S26.**
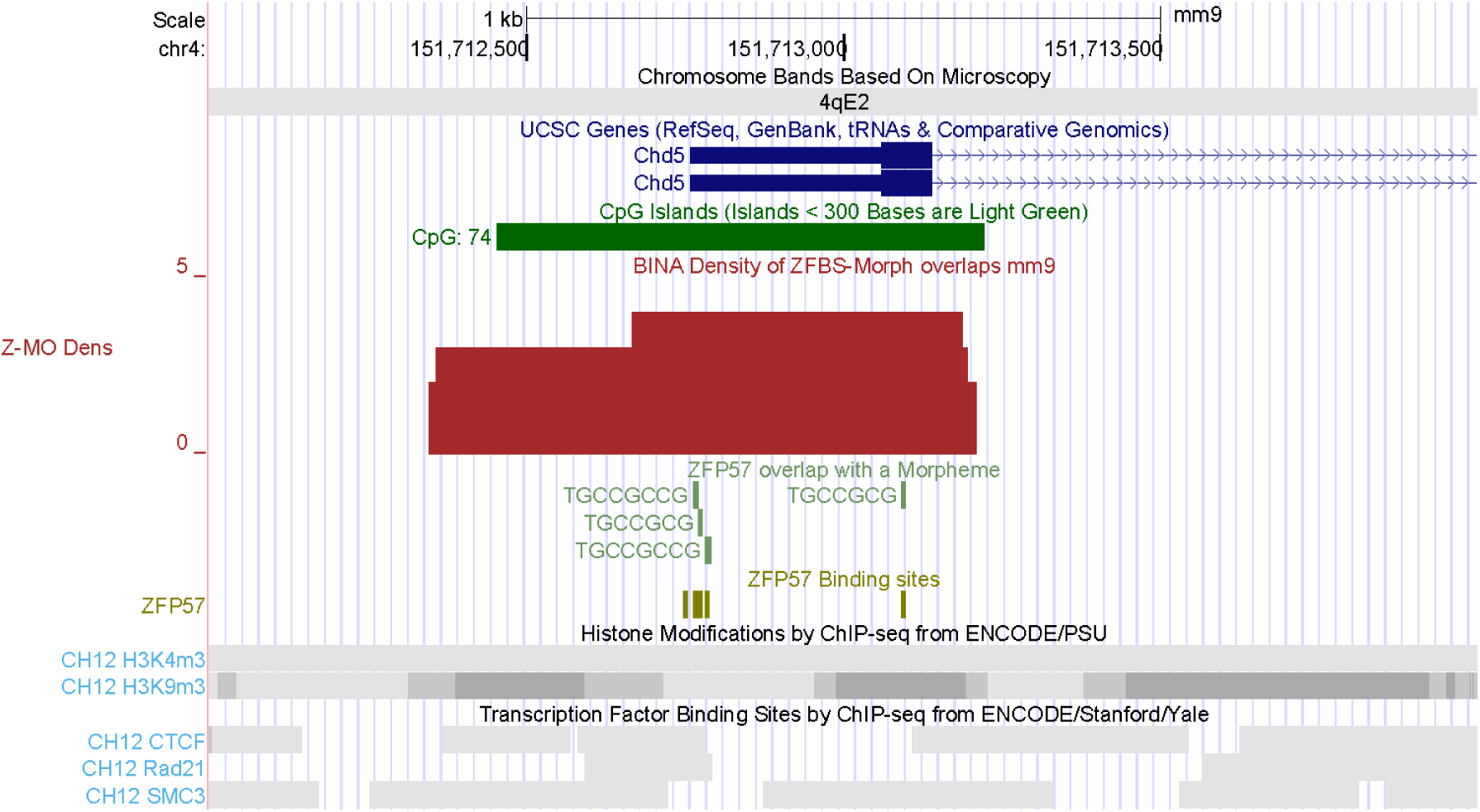
The position of a candidate ICR mapping to *Chd5*. This gene encodes an enzyme that mediates histone-to-protamine replacement and thus impacts the cascade of molecular events underlying chromatin remodeling during spermatogenesis [75].

**Fig. S27.**
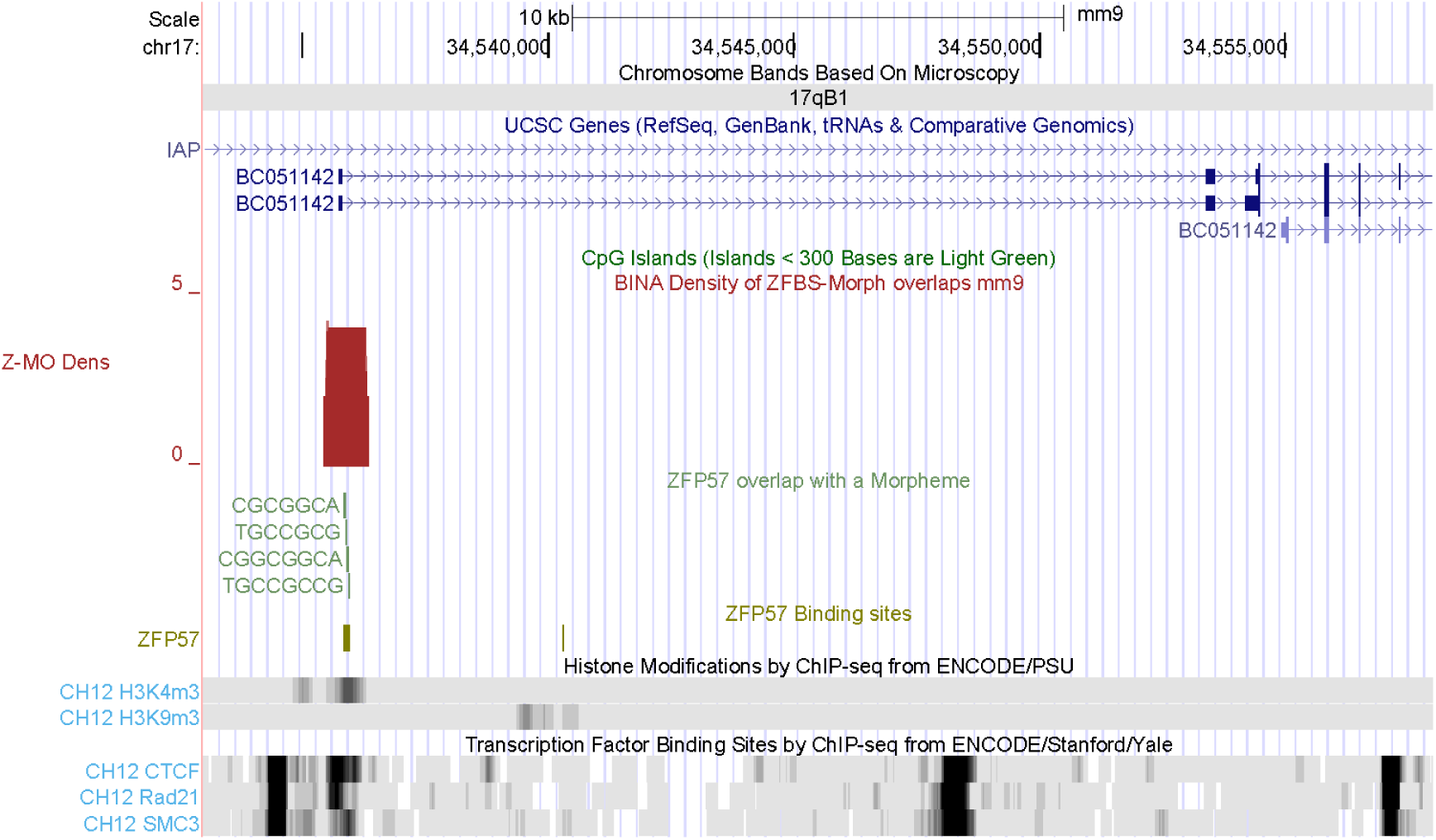
A candidate ICR mapping to longest transcripts of a gene (*BC051142*) expressed in testes. The corresponding density peak maps to a chromatin boundary.

